# Probability weighting arises from boundary repulsions of cognitive noise

**DOI:** 10.1101/2025.09.11.675565

**Authors:** Saurabh Bedi, Gilles de Hollander, Christian C. Ruff

**Author notes:** **Corresponding authors:** Saurabh Bedi; Christian C. Ruff. **Competing Interests:** The authors declare no competing interests.

## Abstract

In both risky choice and perception, people overweight small and underweight large probabilities. While prospect theory models this with a probability weighting function, and Bayesian noisy coding models attribute it to specific encoding functions or priors, we propose a more general account: Probability distortions arise from cognitive noise being repelled by the natural boundaries of probability (0,1). This boundary repulsion occurs in any encoding-decoding system that efficiently encodes, or Bayesian-decodes, bounded quantities, independent of specific priors or encoding functions. Our theory predicts: new, experimentally-induced boundaries should cause additional distortions; increasing cognitive noise should amplify distortions; and boundaries should reduce behavioral variability near them. We confirmed all predictions in three pre-registered experiments spanning risky choice and probability perception. Our findings further suggest that these changes originate largely during decoding. Our work provides a unified explanation for distorted and variable probability judgments, reframing them as consequences of bounded, noisy cognitive inference.

**Significance:** The origin of probability weighting—a central feature of decision-making under risk—remains a longstanding puzzle. Does it arise from processes unique to risk, or does it reflect a more general cognitive mechanism? Here, we show that the classic probability weighting pattern is not domain-specific but instead emerges from a general property of noisy inference over bounded quantities, such as probabilities. Our account formalizes how resource-rational encoding and Bayesian optimal decoding naturally lead to interactions between cognitive noise and the 0–1 bounds of probability, giving rise to systematic distortions. Using pre-registered experimental manipulations across both risky lottery valuation and probability perception, we demonstrate that distortions in probability weighting and estimation are not fixed, intrinsic features, but rather predictable consequences of the interaction between noise and boundaries. This provides a mechanistic account of probability weighting and suggests a unifying explanation for its emergence across different cognitive domains. Similar mechanisms should extend to other naturally or contextually bounded quantities.

## Main

Uncertainty permeates all aspects of our behavior, from everyday decisions to high-stake financial and policy decisions. Human decisions under uncertainty often deviate from normative theoretical models^1^ such as Expected Utility Theory (EUT)^2,3^, which assume that people weigh risky outcomes by their objective probabilities. In reality, people behave as if they distort these probabilities by overweighting small chances and underweighting large ones, a pattern famously described by Prospect Theory’s probability weighting function^4,5^. Yet, while Prospect Theory provides a useful descriptive account, it leaves open the key questions of why probability weighting arises in the first place, and how its underlying cognitive mechanisms may lead to variations in its strength across different choice contexts. Moreover, while Prospect Theory captures average risk attitudes, it offers no account of the systematic variability observed in human decision-making^6,7^.

For these reasons, more recent theories of risky choice have started to incorporate perspectives from models describing how variability in behavior may arise from neural noise inherent in cognition^8^. Inspired by frameworks from neuroscience that model perception as noisy inference^9^, recent models have attempted to explain both probability weighting and behavioral variability as consequences of internal noise in the brain’s process of inferring percepts from sensory information. However, these models often rely on strong and sometimes conflicting assumptions. Some posit that probability weighting arises because probabilities in the environment come from a U-shaped prior distribution with greater density at extremes than intermediate probabilities^10^ and that the brain efficiently encodes probabilities under this prior^11–13^. Others assume the opposite: an inverse-U-shaped prior^14^ for Bayesian decoding^15,16^. These assumptions not only contradict each other but often do not faithfully reflect the priors that would be consistent with the actual probabilities used in the specific experimental context, rendering such inference suboptimal. Yet other models appeal to fixed encoding transformations, such as representing probabilities in log-odds space^17–19^. In these models, the probability weighting function emerges from fixed structural assumptions about how information is transformed during inference. This use of hard-coded priors or non-linear encoding functions has led to the criticism that corresponding Bayesian inference models may have too many degrees of freedom, so that they may in principle capture any type of behavior via highly specific assumptions, without identifying a general underlying mechanism^20–22^.

Here we propose a more general and parsimonious account of the cognitive mechanisms that give rise to probability weighting, which does not have to resort to specific priors or encoding functions. Our account rests on a fundamental property of probabilities: they are naturally *bounded* between 0 and 1. This property, when taken into account either by an efficient encoder or a Bayesian decoder during the noisy cognitive inference process, inevitably leads to truncation of the Bayesian posterior near the boundaries, which systematically repels noisy estimates away from the extremes—a phenomenon we call *cognitive boundary repulsion*. We show that this effect arises robustly across a wide range of prior shapes and encoding functions, and that it can originate independently from both efficient encoding and Bayesian decoding, as long as the quantity that is being inferred is bounded. The general view that probability distortions may relate to truncation is consistent with existing perspectives in economics and perception science. In economics, some studies have fit empirically observed probability weighting by assuming that people calculate expected utilities with random errors truncated for the highest and lowest lottery outcomes^23^. However, our account differs from this in specifying exactly *where* and *why* the truncation occurs, as well as in *what distortions* it predicts. That is, while earlier studies illustrated the descriptive utility of assuming truncation for capturing behavior, our account explains its emergence as a property of the noisy cognitive mechanisms used to infer any bounded quantities. Moreover, our account makes novel, counterintuitive experimental predictions that we test empirically. In perception, Bayesian decoding models have explained related perceptual biases with bounded (often uniform) priors that are consistent with experimentally imposed stimulus ranges^24–26^, showing that uniform bounded priors can generate systematic distortions even for contextually bounded stimuli (not naturally bounded like probabilities). Our account goes further in demonstrating that these effects can arise not only during decoding but also at encoding, across diverse bounded prior distributions and encoding functions. While specific priors or encoding functions may influence features such as the crossover point—where overweighting shifts to underweighting—they are not required for the core pattern to emerge, offering a general explanation for the ubiquity of probability distortions.

In our account of cognitive boundary repulsions, distortions can occur both at the efficient encoding and Bayesian decoding stages. However, these two potential sources of distortion would lead to different behavioral signatures that can be empirically disentangled. At the encoding stage, we show that boundary repulsions arise in the likelihood when the brain efficiently adapts its limited resources to maximize mutual information between bounded quantities and their biophysically constrained, bounded representations^12,27–29^. Such efficient encoding adaptation^30,31^ causes range dependence of neural representations and behavior, as widely observed across sensory domains^32–34^, perceptual choice^35^, and value-based decision-making^36–38^, and formalized in process-level theories such as Range–Frequency Theory^39^ and Decision-by-Sampling^40^, both of which have efficient-coding interpretations^41^. Here, we show that if there is adaptive efficient coding for bounded quantities, it necessarily produces boundary repulsions in cognitive noise that scale with the input range. As the range of inputs increases, the same boundary repulsion within the representational space maps onto a larger bounded quantity space—producing stronger distortions and greater variability in behavior. However, when boundary repulsions arise only at the time of decoding, the encoded representations remain fixed and do not rescale with changes in the bounded range. Still, Bayesian decoding with a bounded prior will truncate the posterior distribution, but without rescaling. As a result, the distortions and behavioral variability remain invariant to the size of the input range. Thus, while both encoding and decoding can independently produce boundary repulsion effects, they generate distinct patterns: boundary repulsion arising during encoding scales with the size of the range, whereas boundary repulsion arising during decoding does not.

Most accounts of probability weighting are inherently context-independent. They assume that distortions reflect something intrinsically special about small and large probabilities^4,5,42^, the highest and lowest outcomes of lotteries^23^, or an outcome sampling mechanism for inferring probabilities^43,44^. Some context-dependent theories rely on long term statistics, such as assuming that small and large probabilities occur more often in the environment^10,40^. Bayesian models can flexibly account for context-dependent distortions by varying assumed prior shapes, but most existing accounts attribute reduced variability near the extremes to fixed transformations like log-odds encoding^17,19^, which do not predict changes in variance near the boundaries across contexts. By contrast, our account explains probability weighting as a natural consequence of noisy inference over bounded quantities. This mechanism uniquely predicts a joint behavioral signature that is not predicted by the other existing theories: when new boundaries are introduced, new distortions should arise, and response variability should decrease near those boundaries. These effects should appear in both simple perceptual judgments and higher-order lottery valuations. We tested these predictions in three pre-registered experiments across perceptual and value-based tasks, in which we manipulated cognitive noise levels and introduced explicit contextual boundaries, via instructions informing participants of the range of probabilities they could encounter. Consistent with our predictions, we found that (1) higher cognitive noise amplified probability distortions, (2) newly introduced boundaries induced additional probability distortions, and (3) variance in estimates systematically decreased near both natural and contextually induced boundaries. Moreover, both distortion and variability remained largely range-independent—consistent with the idea that the boundary repulsion arises from the Bayesian decoding stage, rather than at the encoding stage. Our account thereby grounds the origins of probability weighting in noisy inference of bounded quantities and reveals principles likely to generalize across other domains of cognition involving bounded quantities.

## Results

### Boundary repulsions in noisy inference of bounded quantities

Many cognitive tasks, ranging from perceptual estimation to valuation and decision making, involve stimuli that lie within a bounded range. One example is probabilities that naturally lie between 0 and 1. Critically, such tasks require decision-makers to represent variables within a noisy neural system, which inevitably introduces variability and systematic bias into these representations^45^. We model this phenomenon using the well-established general framework of noisy encoding followed by decoding^9,13^, in which a stimulus *b* is encoded into a noisy internal representation *η* via the encoding function *F*(*b*) plus corrupting noise ϵ, and subsequently decoded to generate an inferred estimate 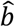 (Fig. 1a). This estimate is either reported directly in perceptual judgment tasks or is used as input to subsequent valuation/value comparison in risky decision-making tasks. When applied to any bounded quantity, irrespective of its assumed distribution as captured by a wide variety of prior shapes, this process ultimately leads to a truncated posterior that is skewed away from the boundaries whenever inference involves either efficient coding or Bayesian decoding or both (Fig. 1a; see Methods). We call this emergent inward asymmetry of the posterior distribution *cognitive boundary repulsions* (Fig.1f). These repulsions lead to the characteristic *probability weighting function* (Fig. 1g), where small probabilities are overweighted and large probabilities are underweighted due to overestimation of the mean posterior estimate above the lower boundary and underestimation below the upper boundary (Fig.1h). Additionally, the repulsions predict reduced variability of the estimate near boundaries (Fig.1i).

**Fig. 1.**
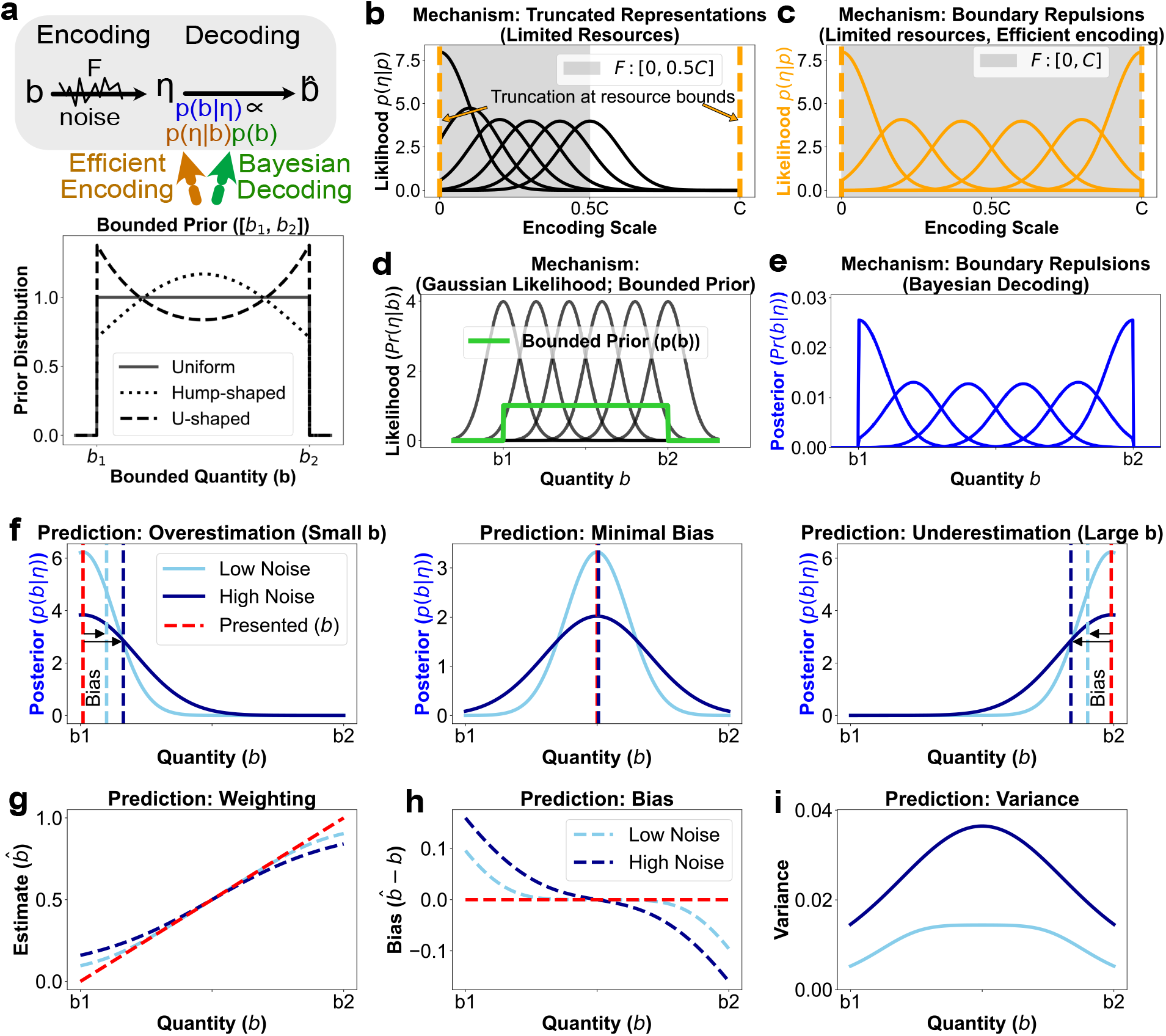
Cognitive boundary repulsions in noisy inference of bounded quantities. **a)** Bounded quantities (e.g., probabilities between 0 and 1) are encoded into noisy internal representations η and then decoded to form an estimate 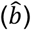. This inference process can be influenced by the bounded nature of the stimulus either at the *encoding stage* via efficient allocation of limited representational resources, or at the *decoding stage*, via Bayesian decoding with a bounded prior. Bottom: Similar general effects arise across diverse prior shapes (uniform, U-shaped, inverse-U-shaped), emphasizing that boundary repulsions are not dependent on specific prior shapes. **(b)–(f)** Schematic illustrations of the mechanisms captured, and predictions derived from, our account. **(b)** When encoding is limited to a finite representational capacity *η* ∈ [0, C], encoding noise becomes truncated near representational bounds. **(c)** If encoding further optimizes mutual information between the input and representations, the allocation of bounded representational capacity across the bounded input range induces inward asymmetry in the likelihood function *p*(*η*∣*b*), even before decoding. **(d)** Even with symmetric likelihoods (e.g. from unconstrained or non-adaptive encoding), applying bounded priors affect the posterior through Bayes’ rule. **(e)** This leads to systematic inward skew in the posterior near boundaries, even when encoding stage did not induce asymmetries in the likelihood. **(f)** Both efficient encoding and Bayesian decoding mechanisms independently produce truncated, inward-skewed posteriors that repel inferred estimates away from boundaries of inputs (e.g., 0 and 1 for probabilities), giving rise to cognitive boundary repulsions. **(g)** Posterior means are biased away from boundaries, producing the characteristic shape of probability weighting (overweighting of small probabilities and underweighting large ones). **(h)** The magnitude of distortion increases with noise. **(i)** Variance of estimates is reduced near boundaries, another signature of cognitive boundary repulsions.

First, we formally derive (in Methods) that, given the framework of noisy encoding and decoding, and under some weak assumptions of optimality, cognitive boundary repulsions *have to* occur for at least two reasons (see Methods): (1) when *encoding* is efficient in the sense that it maximizes mutual information^13,46^ between the bounded inputs and a bounded representational space, likelihood (noise) distributions will be skewed inwards, and (2) when *Bayesian decoding* is performed with any (objective) bounded prior, mean posteriors will be biased inwards as well, irrespective of the shape of this prior. In other words, at the *encoding stage*, if internal representations are limited *η* ∈ [0, *C*] then this inevitably leads to truncated likelihoods (Fig.1b) as there is a finite support for encoding. Our derivation shows that if these bounded representations efficiently optimize to maximize mutual information about the bounded quantity *b* ∈ [*b*_1, *b*_2], then the encoding function must span the dynamic range of available resources in a way that leads to inward skew in encoding noise and likelihood, pushed away from the boundaries (Fig.1c). In contrast, at the *decoding stage*, even if encoding is unbounded or non-adaptive (Fig.1d), boundary repulsions arise purely from applying Bayes’ rule with a *bounded prior*. The prior truncates the posterior at the boundaries, causing systematic inward skew of the mean posteriors —even when the likelihood is symmetric and not truncated (Fig.1e). We validated both encoding- and decoding-based repulsions in simulations across diverse priors and encoding functions (see Supplementary Figs. 1-3). For numerical simulations we implemented the cumulative distribution function (CDF) transform of the prior, a redundancy-reducing code for efficient coding^12^ (see Methods and Supplementary for details). In line with our account and its predictions, encoding-based effects simulated across different prior shapes reproduced the characteristic probability weighting pattern (Supplementary Fig. 1). Similarly, the characteristic probability weighting pattern also emerged also for decoding-based effects under various prior shapes and encoding functions (see Supplementary Figs. 2 and 3). Together, our theoretical derivations and numerical simulation results show that characteristic probability weighting patterns can arise solely due to cognitive boundary repulsions, as consequence of noisy inference over bounded quantities, at both the encoding and decoding stage.

Although cognitive boundary repulsions can occur both at encoding and decoding stage, we also show that they produce *dissociable behavioral signatures* (see Methods). Encoding-based repulsions scale with the range of the bounded inputs. As the stimulus range widens, a fixed-capacity representational space must be stretched to cover a larger input interval. As a result, the same internal noise maps onto a broader range of inputs, amplifying both distortions and variability in the inferred estimates. Even *partial adaptation*, where encoding only adjusts partially to the input range produces *range adaptation* of cognitive boundary repulsions. This result extends the general range adaptation of neural representations and behavior observed for efficient adaptation in various contexts^30–39,47^. In contrast, when boundary repulsions arise solely during the decoding stage, i.e., when encoding is completely non-adaptive, the representational space remains unchanged. As a result, the magnitude of boundary repulsions of estimates remains invariant to input range, because internal noise continues to map to the same inputs. We also demonstrate this theoretical dissociation in simulations (Fig.7, left column), using the CDF approximation of efficient encoding and a uniform prior, where the ultimate percept is defined as the mean posterior.

To test the predictions of our cognitive boundary repulsions account, we designed a series of experiments that independently manipulated its two key variables: the amount of cognitive noise and the presence of contextual boundaries. Across tasks involving risky decision-making and purely perceptual probability estimation, we varied the complexity of fractions that represented probabilities (to modulate noise) and explicitly instructed participants about the range of probabilities they would encounter in each block (to introduce contextual boundaries). Crucially, we also distinguished these cognitive effects from a separate class of boundary effects: Mechanical response truncation in errors. In many tasks, responses are constrained by fixed bounds (e.g., sliders or number ranges), such that errors in estimates exceeding the limits are forcibly clipped. When participant-level biases differ, such mechanical truncation can produce apparent group-level truncation in errors^48^ that may look like cognitive boundary repulsions. To isolate cognitive boundary repulsions from potential non-cognitive, response-scale artifacts, we designed our boundary manipulations so that they did not alter the response scale itself. Therefore, contextual boundary manipulations in our experiment were executed without altering the output scale for lottery valuations and probability estimate responses, thereby eliminating the possibility of non-cognitive, bounded output range effects.

### Testing core predictions of the cognitive boundary repulsion account

Our account proposes that probability weighting arises not from fixed encoding transformations or prior shapes, but from a general computational principle: when bounded quantities are inferred under noise, they give rise to systematic repulsions of the posterior away from the boundaries. These cognitive boundary repulsions, emerging naturally at boundaries of 0 and 1 for probabilities, produce biases in estimated probabilities that propagate and produce the characteristic probability weighting observed in risky valuation tasks as described in Prospect Theory. However, they are accompanied by additional, diagnostic signatures in variability that reflect the underlying boundary repulsions in noise due to noisy inference of bounded quantities.

Our cognitive boundary repulsion account makes concrete, testable predictions. First, increasing noise should make probability weighting more extreme. Second, introducing novel boundaries into a stimulus space should produce repulsive biases like those induced by natural boundaries. Third, adding these boundaries should result in diminished variance patterns near the new boundaries. Finally, this mechanism and its predictions should be domain-general: it should apply across tasks that require probabilistic inference, whether probabilities are estimated directly for perceptual report or used indirectly for valuation or value-based choice.

To test these predictions empirically, we conducted three pre-registered experiments (https://osf.io/8sykp, https://osf.io/cfgbr, https://osf.io/vzdsq) across two distinct cognitive domains. Experiments 1 and 2 were risky lottery valuation tasks (Fig. 2a) in which participants reported certainty equivalents (CEs), indicating the minimum sure payoff they would accept to give up the opportunity to take part in risky lotteries paying 50 ECU (Experimental Currency Units) at varying probabilities. Experiment 3 was a perceptual judgment task (Fig. 2d) in which participants had to estimate probabilities corresponding to fractions of different complexity. In all experiments, we manipulated two key components of the inference process within-subject: *cognitive noise*, by altering the complexity of fraction formats used to present probabilities (e.g., simple fractions like 35/100 vs. complex ones like 2334/8049) (Fig. 2c,f); and *contextual boundaries*, introduced via block-wise explicit instructions about the probability range participants would encounter (e.g., [0–0.5], [0.5–1], or [0.34–0.66]) (Fig. 2b, e). Probabilities were sampled uniformly within each range, and participants were explicitly informed of the probability range and the uniform sampling before each block, creating contextual boundaries. Because our boundary manipulations depended on participants’ understanding of these induced ranges, comprehension questions were asked at the start of every block to ensure participants correctly knew the range of probabilities they were about to judge or decide on in the upcoming block.

**Fig. 2.**
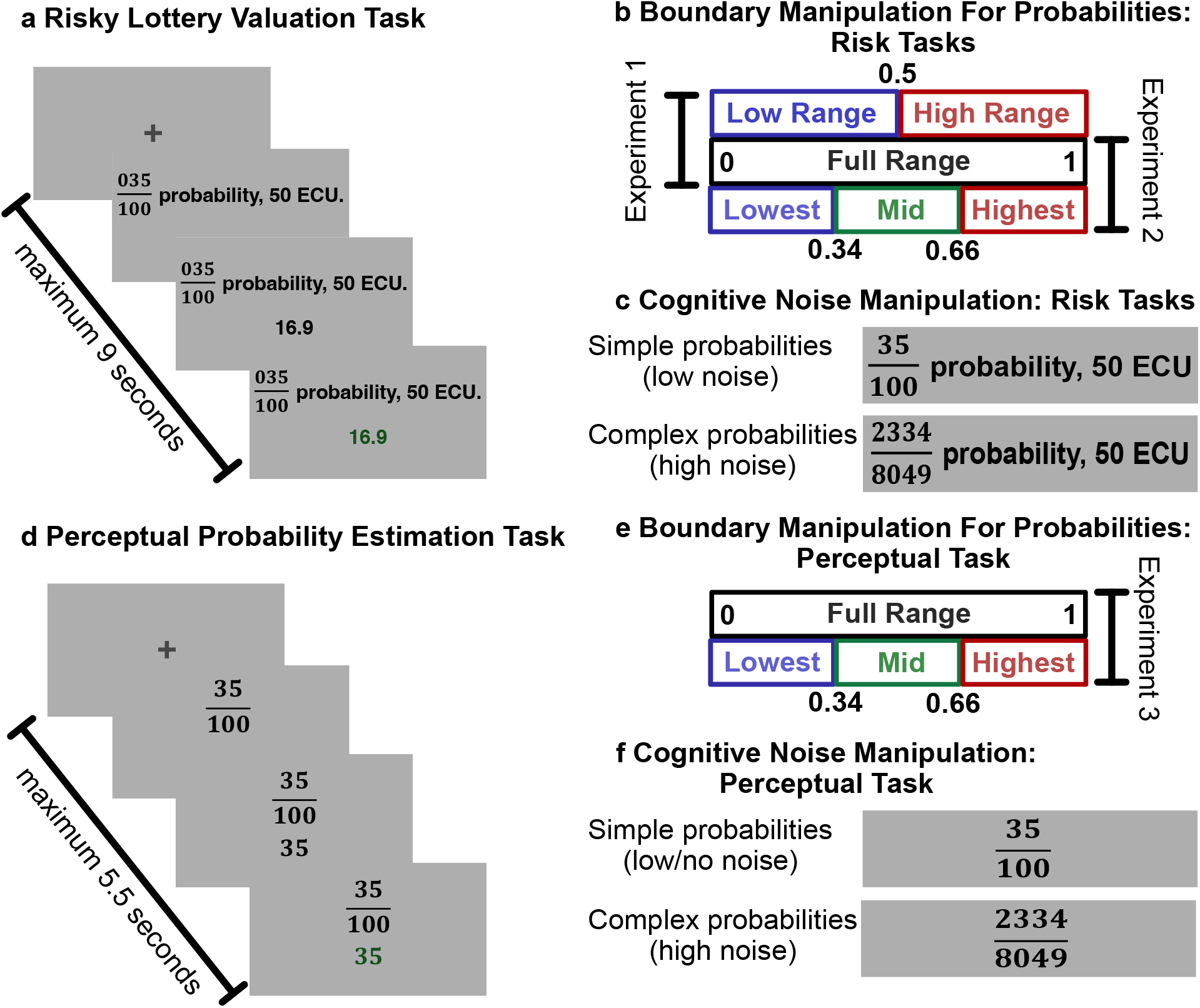
Experiment design: (a) Risky Lottery Valuation Task (Experiments 1 & 2): On each trial, participants valued risky lotteries, reporting their certainty equivalent (CE) for a lottery that paid 50 ECU with some probability and 0 otherwise. The entered CE turned green once the participants had confirmed their entry. (**b) Boundary Manipulations in the Risk Task:** Before each block, participants were explicitly informed of the probability range they would encounter, inducing contextual probability boundaries. Experiment 1 introduced a single boundary at 0.5, while Experiment 2 added two boundaries at 0.34 and 0.66. Probabilities in each block were sampled uniformly and never exceeded the stated range. **(c) Cognitive Noise Manipulation in the Risk Task:** All probabilities were presented as fractions, with difficulty manipulated across blocks. Low-noise blocks used simple fractions with a fixed denominator of 100, while high-noise blocks used complex four-digit fractions to increase difficulty in inferring them. **(d) Perceptual Judgment Task (Experiment 3):** Participants viewed probabilities as fractions on each trial and estimated the corresponding percentage as accurately as possible. **(e) Boundary Manipulations in the Perceptual Task:** Trials were grouped into blocks based on four explicitly stated probability ranges: 0-1, 0-0.33, 0.33-0.65, and 0.67-1, introducing varying contextual boundaries. (**f) Cognitive Noise Manipulation in the Perceptual Task:** Noise was manipulated via fraction complexity. In low/no noise blocks, participants only had to enter the numerator (fixed denominator of 100); in high noise blocks, both numerator and denominator were four-digit values requiring full fraction evaluation, thus increasing cognitive difficulty.

This experimental setup enabled us to evaluate core predictions of the cognitive boundary repulsion account; each directly derived from the model’s core claim: noisy inference of bounded quantities leads to repulsive biases and reductions in variance near the boundaries.

### Amplifying probability distortions by increasing cognitive noise

The first prediction of our account is that increasing cognitive noise should amplify repulsions at the natural bounds of probability (0 and 1), thereby strengthening both the overweighting of small probabilities and underweighting of large probabilities. This follows directly from the account’s central assumptions that boundaries repel away noise, so that higher noise should lead to a strengthening of this repulsion at both upper and lower boundaries. To make our predictions more concrete, we ran simulations of our experiment using the noisy inference model with bounded priors during decoding. The results of these simulations and their predictions are plotted next to empirical data (Fig. 3a, e). In the simulations, decision-makers use linear encoding of probabilities, followed by Bayesian decoding with a uniform bounded prior reflecting each experiment’s probability range, and two levels of internal noise (σ = 0.02 and σ = 0.06). To manipulate the noise experimentally, we manipulated fraction complexity as a proxy for cognitive noise. As predicted, participants’ lottery valuations (Experiments 1 & 2; Fig. 3b, c, f, g) and probability estimates (Experiment 3; Fig. 3d, h) showed significantly stronger distortions near the natural boundaries under high cognitive noise. All effects were assessed using a pre-registered mixed-effects model (see Methods) analyzing the within-subject effects of noise (fraction complexity) on the reported certainty equivalent/probability, in preregistered probability bins near the natural boundaries (grey shaded regions in Fig. 3). In all three experiments, we observed the expected within-subject amplification of overweighting/overestimation for small probabilities (all betas > 0, all p < 0.001, see Supplementary Tables 1.1.1 – 1.1.3), and underweighting/underestimation of large probabilities (all betas < 0, all p < 0.001, see Supplementary Tables 1.2.1 – 1.2.3). This confirms that probability distortions indeed increase with higher cognitive noise, an effect that is not predicted by existing models of probability weighting such as Prospect Theory that assume a fixed functional form^4,5,42^ and do not incorporate noise-dependent mechanisms. On the other hand, existing accounts relying on noise^14,17,19,47^ or error based mechanisms^23,43,44,49^ can explain these results.

**Fig. 3.**
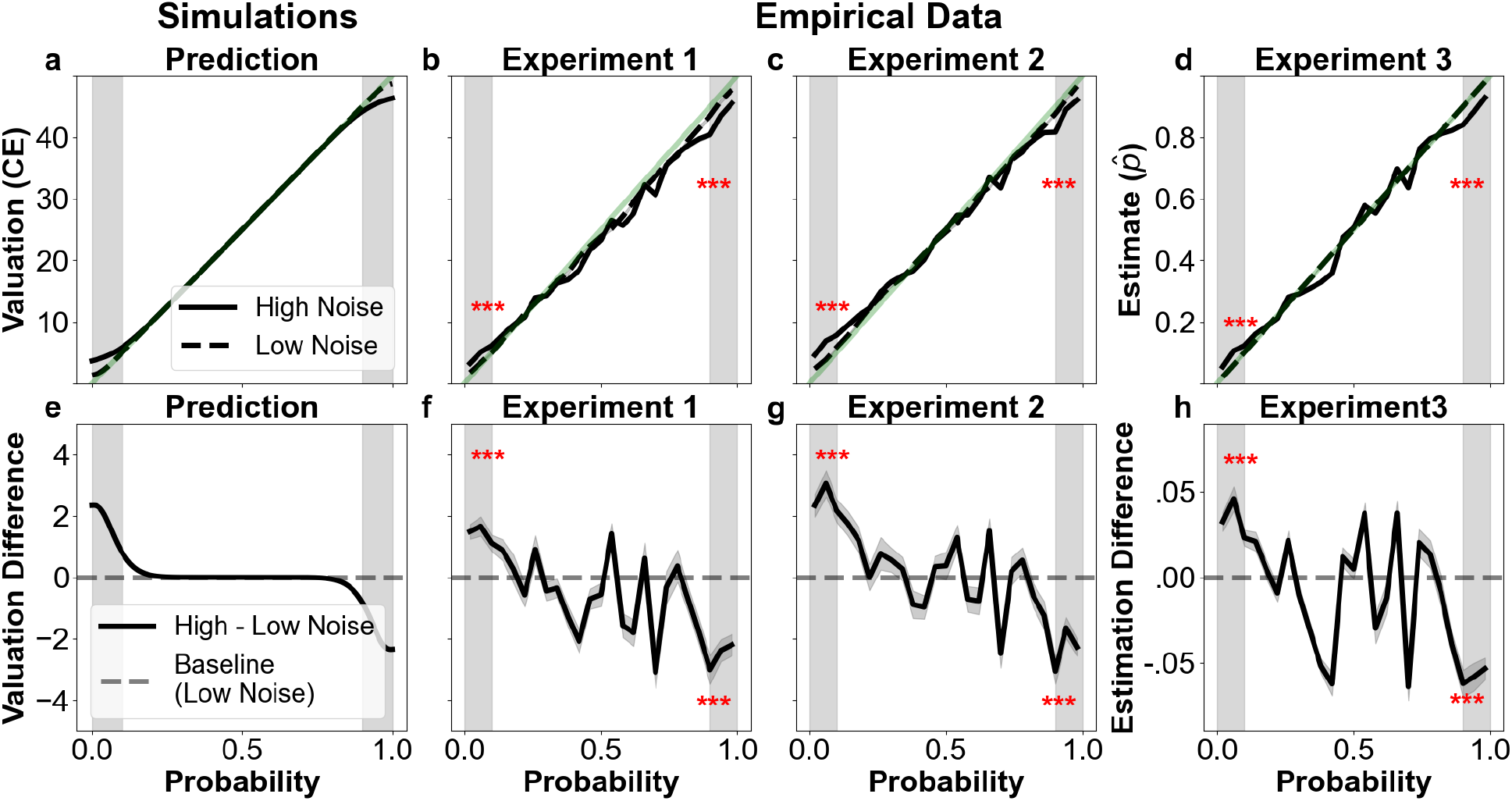
Effect of cognitive noise on probability distortions in risk and perception: **(a, e) Model predictions of noise-dependent distortions**. Simulations from the cognitive boundary repulsion model with linear encoding and a uniform bounded prior (see Methods). Certainty equivalents (CEs) were computed by multiplying the mean of noisy probability estimates with the lottery value (50 ECU). **(a)** Predicted CEs under low (σ = 0.02; dashed line) and high (σ = 0.06; solid line) noise parameters. Higher noise amplifies boundary repulsions, leading to stronger deviations from the expected value line (green). **(e)** Predicted difference in CEs between high and low noise conditions as a function of underlying probability. **(b, f) Empirical results from Experiment 1 (risky lottery valuation task). (b)** Average CEs across probabilities for simple and complex probabilities. **(f)** Difference in lottery valuation between complex and simple probability conditions. As predicted, high noise (complexity) produces stronger probability weighting. Black lines indicate mean valuations/valuation differences; grey bands around the mean indicate ±1 s.e.m. (standard errors of the mean); red asterisks denote significant effects (p < 0.001) in pre-registered bins (grey shading on left and right). **(c, g) Replication in Experiment 2**. Data confirm the same noise dependent distortion patterns in an independent sample. **(d, h) Generalization to perceptual experiment (Experiment 3)**. Participants estimated numeric probabilities from fraction inputs. Results show systematic overestimation of small probabilities and underestimation of large probabilities under high noise. These perceptual distortions mirror the valuation distortions predicted by the model, supporting the domain-generality of the cognitive boundary repulsion account.

### Inducing new boundaries creates new probability distortion patterns

A central prediction of the cognitive boundary repulsion account is that repulsions are not exclusive to natural bounds (0 and 1) but arise for any bounded quantity. Therefore, *experimentally induced boundaries* should induce novel distortions that resemble those found at natural boundaries. Additionally, these new distortions should become stronger under increased cognitive noise. Crucially, this prediction fundamentally departs from existing accounts, as they attribute the classical distortion patterns to something inherently special in the way small and large probabilities are weighed, encoded, or decoded. In our account, small and large probabilities appear special only because of the natural boundedness of probabilities. By altering these boundaries, other probabilities should acquire this special status as well, systematically altering the characteristic distortion pattern.

We tested this prediction by manipulating contextual probability boundaries across three pre-registered experiments. In Experiment 1 (lottery valuation), we introduced a single additional boundary at 0.5. In Experiments 2 (lottery valuation) and 3 (probability estimation), we introduced two new boundaries at 0.34 and 0.66 to replicate the effects of experiment 1 with additional boundaries and across tasks. In all cases, participants completed blocks in which probabilities were drawn uniformly either from the full range (0,1) or from restricted subranges defined by new boundaries. Since all probabilities and response formats were held constant, any observed changes must result from altered probability inference due to newly imposed boundaries, providing a strong test of our account.

We conducted two pre-registered analyses using the same mixed-effects model that was applied to test our first hypothesis (see Fig. 3 and Methods). Again, all analyses were done on pre-registered probability bins (grey bins in Figs. 4 and 5). First, we tested an *interaction effect* (Fig. 4) to assess whether introducing contextual boundaries, combined with increased probability complexity (cognitive noise), induced new distortions. Second, we tested a *main effect* (Fig. 5) examining whether boundaries alone, under high noise, reshape estimation and valuation for *identical* probabilities/lotteries within-subjects. The simulations plotted next to empirical data in Figs. 4 and 5 used the same model parameters as in Fig. 3: linear encoding and Bayesian decoding with a uniform bounded prior reflecting each experiment’s probability range, and two levels of internal noise (σ = 0.02 and σ = 0.06). For *Experiment 1*, the interaction effect was visualized by simulating predicted valuations for the full-range, low-noise condition and for the half-range, high-noise conditions (Fig. 4a), and plotting their difference to capture the predicted interaction effect (Fig. 4c). For *Experiments 2 and 3*, which included two boundaries, the interaction effect is visualized by plotting simulated differences between full-range low noise and restricted-range high-noise conditions of valuation and estimation respectively (Fig. 4e, g). The resulting plots predict valuation/estimation dips below and jumps above each added boundary, due to the interaction of cognitive noise and boundaries. For the main effect, we simulated high-noise conditions with and without contextual boundaries (Fig. 5), holding noise constant to isolate the influence of boundaries alone. The resulting difference plots (Fig. 5c, e, g) show how boundary structure alone alters valuation and estimation.

**Fig. 4.**
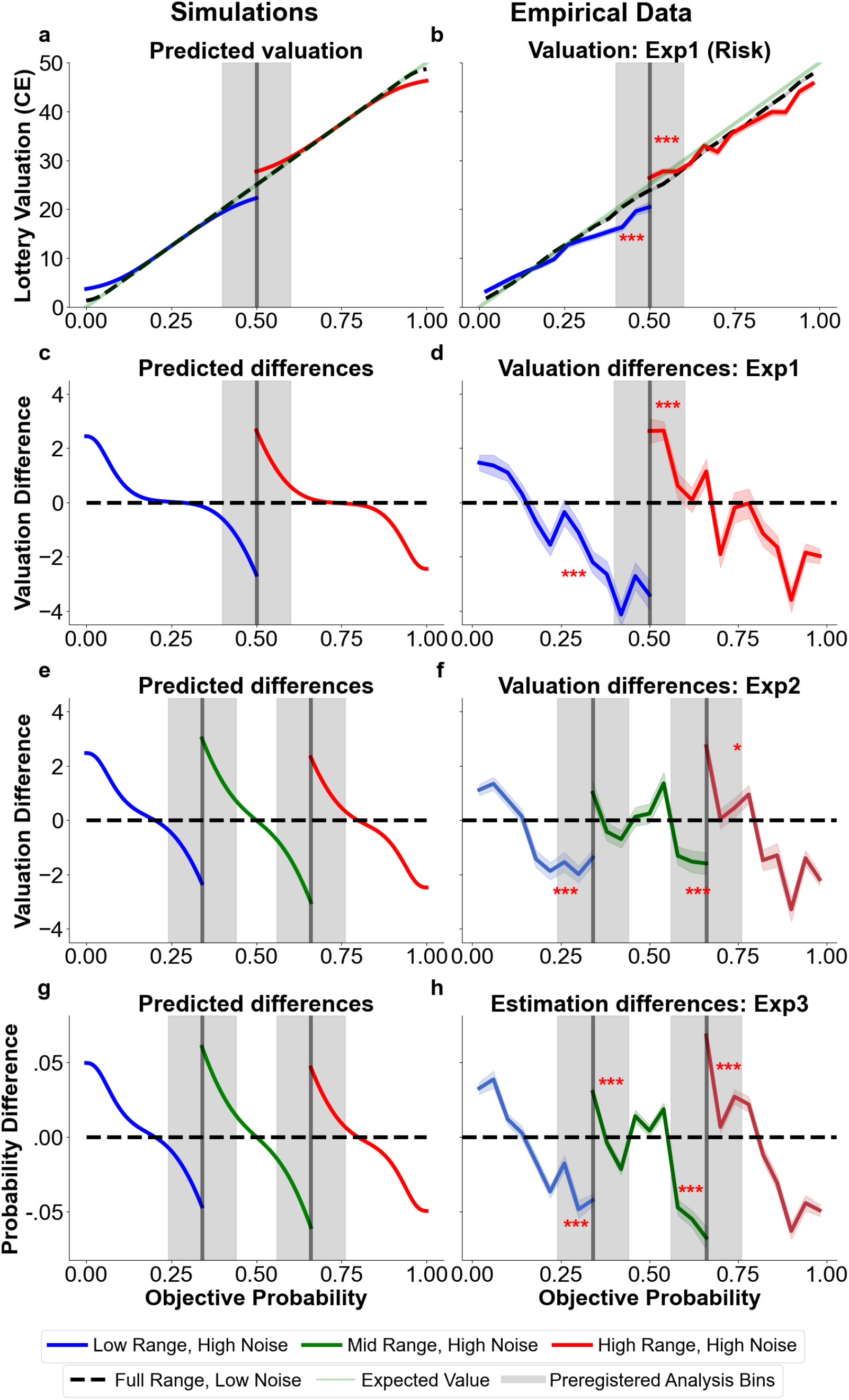
Interaction effect of induced boundaries and cognitive noise: **(a, c, e, g)** Model predictions from the cognitive boundary repulsion account, assuming linear encoding and Bayesian decoding with a uniform prior bounded within the experimental block ranges (see Methods). Simulations illustrate predicted distortions when comparing high-noise (σ = 0.06) and low-noise (σ = 0.02) conditions across different contextual boundaries. **(a)** Predicted certainty equivalents (CEs) in a lottery task with full-range probabilities (black dotted line) versus restricted-range probabilities with an added boundary at 0.5 (red/blue lines), showing the emergence of a double weighting pattern under high noise. **(c, e, g)** Simulated interaction effects—computed as the difference in predicted valuations or estimations between high-noise restricted-range and low-noise full-range conditions—demonstrate how boundary repulsions emerge symmetrically around induced boundaries at 0.5 (c) and at 0.34 and 0.66 (e, g). **(b, d, f, h)** Empirical results validating these predictions across three experiments. **(b)** In Experiment 1 (lottery valuation), participants show distorted valuations consistent with a double weighting pattern when a boundary at 0.5 is introduced under high noise. **(d)** The corresponding interaction effect is visualized as valuation differences between conditions, with increases above and decreases below the induced boundary. **(f)** In Experiment 2 (lottery valuation), introducing two boundaries at 0.34 and 0.66 under high noise produces a triple weighting pattern, with valuations increasing above and decreasing below each boundary. **(h)** In Experiment 3 (perceptual probability estimation), the same manipulation leads to equivalent distortions in subjective probability estimates, demonstrating that boundary repulsion generalizes beyond valuation to perceptual inference. Grey shaded areas mark pre-registered probability bins used for statistical testing. Black lines show means; grey bands indicate ±1 standard error of the mean (s.e.m.); single and triple red asterisks denote significant effects (*p* < 0.05 and *p* < 0.001, respectively) within pre-registered bins.

**Fig. 5.**
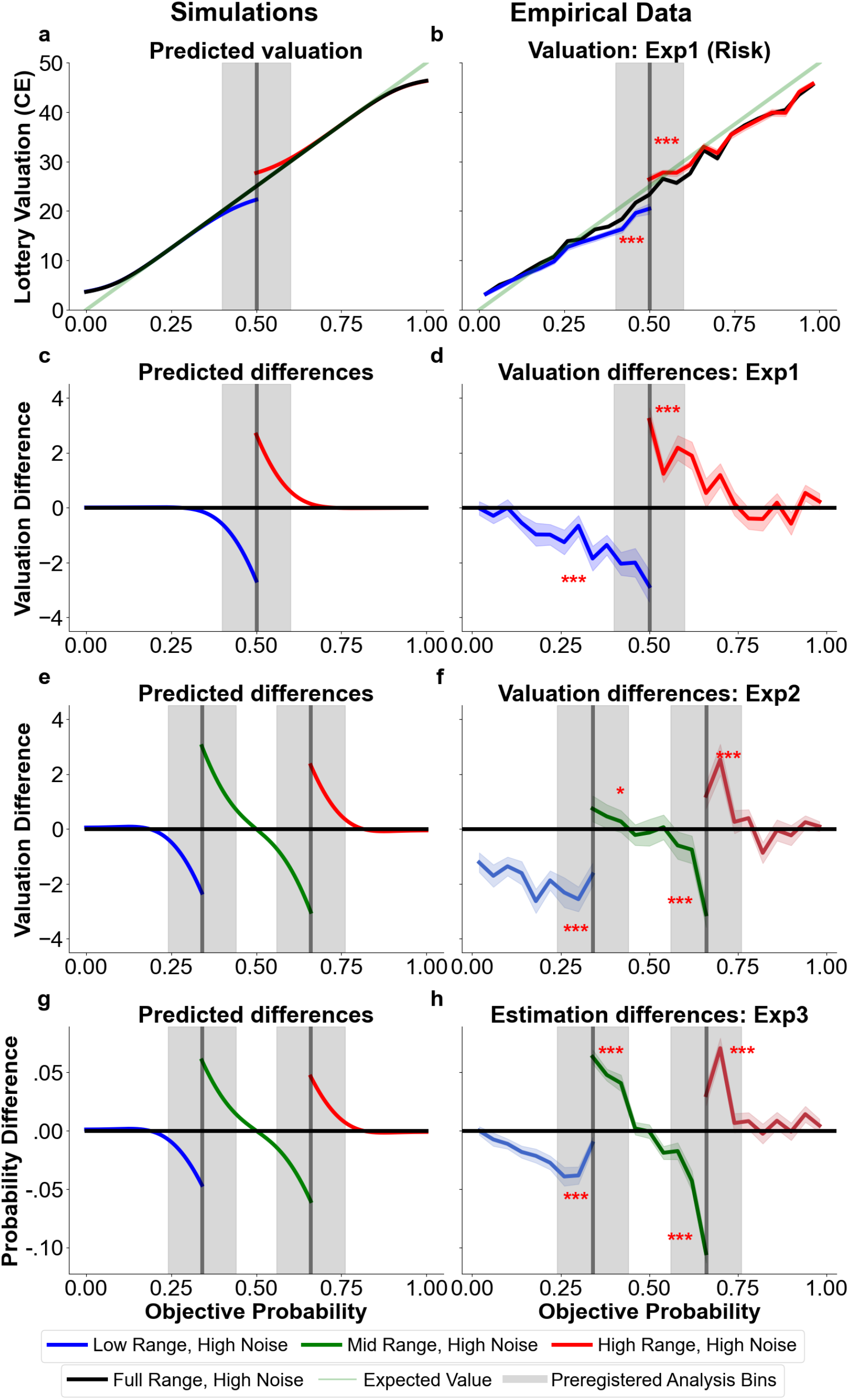
Main effect of induced boundaries on probability distortions for high cognitive noise: **(a, c, e, g)** Model predictions from the cognitive boundary repulsion account under high internal noise (σ = 0.06), comparing restricted-range blocks with added boundaries to full-range blocks without boundaries. Simulations assume linear encoding and Bayesian decoding with a uniform prior bounded by the experimental block ranges (see Methods). **(a)** Predicted certainty equivalents (CEs) in a lottery task show a single weighting function in the full-range condition (black), but a double weighting pattern when a boundary is introduced at 0.5 (red/blue), consistent with boundary repulsion. **(c, e, g)** Simulated main effects—computed as the difference in predicted valuations or estimations between restricted-range and full-range conditions—demonstrate how boundary-induced distortions emerge around contextual boundaries at 0.5 (c) and at 0.34 and 0.66 (e, g), even when noise is held constant. **(b, d, f, h)** Empirical results confirm these predictions across three experiments. **(b)** In Experiment 1 (lottery valuation), introducing a boundary at 0.5 under high noise produces a double distortion pattern in valuations. **(d)** This main effect is visualized as systematic shifts in valuations above and below the induced boundary. **(f)** In Experiment 2 (lottery valuation), adding boundaries at 0.34 and 0.66 produces a triple weighting pattern, with systematic valuation shifts above and below each boundary. **(h)** In Experiment 3 (perceptual probability estimation), the same boundary manipulation produces equivalent distortions in subjective probability estimates, demonstrating that boundary repulsion generalizes across domains. Grey shaded areas mark pre-registered probability bins used for statistical testing. Black lines show mean responses; grey bands represent ±1 standard error of the mean (s.e.m.); single and triple red asterisks denote significant effects (*p* < 0.05 and *p* < 0.001, respectively) within pre-registered bins.

In *Experiment 1*, adding a single contextual boundary at 0.5 during risky lottery valuation significantly altered the classic probability weighting pattern. As predicted, we observed both an *interaction effect* (Fig. 4b, d) and a *main effect* (Fig. 5b, d), showing that the addition of the new artificial boundary, together with cognitive noise, was sufficient to distort valuations for identical lotteries. Specifically, the classic probability weighting pattern was transformed into the predicted double-distorted weighting pattern, with the expected significant increase in valuations above the contextually induced boundary of 0.5 (interaction: beta > 0, *p* < 0.001; see Supplementary Table 2.1.1; main effect: beta > 0, *p* < 0.001; see Supplementary Table 3.1.1) and the significant decrease in valuations below the contextually induced boundary (interaction: beta < 0, *p* < 0.001; see Supplementary Table 2.2.1; main effect: beta < 0, *p* < 0.001; see Supplementary Table 3.2.1). In *Experiment 2*, we replicated this boundary-induced distortion using *two additional boundaries* (at 0.34 and 0.66), again confirmed through both the interaction (Fig. 4f) and main effects (Fig. 5f). This showed that the phenomenon generalizes beyond a single boundary, and can produce even a triple-distorted weighting pattern when two additional contextual boundaries are induced. The interaction effect confirmed the expected increased valuations above both induced boundaries, although it reached significance only in one of the two bins (both betas > 0; *p* < 0.05 for 1 bin; see supplementary tables 2.1.2 and 2.1.3). The main effect confirmed significantly increased valuations in both pre-registered bins (both betas > 0, with significance in one bin reaching *p* < 0.05, and the other *p* < 0.001; see supplementary tables 3.1.2 and 3.1.3), and both the interaction effect and the main effect confirmed significantly decreased valuations in both pre-registered bins (all betas < 0, all *p* < 0.001; see supplementary tables 2.2.2, 2.2.3, 3.2.2, 3.2.3). Finally, in *Experiment 3*, we replicated these effects in the domain of *perceptual probability estimation*, showing that the same boundary and noise manipulations produced equivalent distortions for purely perceptual judgments outside the domain of risky decision-making (Fig. 4h, Fig. 5h). Also in this task, both the interaction and the main effects confirmed a significant increase in estimated probabilities in both pre-registered bins (all betas > 0, all *p* < 0.001; see Supplementary Tables 2.1.4, 2.1.5, 3.1.4, 3.1.5) and a significant decrease in estimated probabilities in both pre-registered bins (all betas < 0, all *p* < 0.001; see Supplementary Tables 2.2.4, 2.2.5, 3.2.4, 3.2.5). Together, these results demonstrate that probability weighting is not fixed but is systematically and flexibly shaped by the structure of contextual boundaries and cognitive noise, supporting the boundary repulsion account across both valuation and perceptual tasks.

These results confirm that multiple contextual boundaries produce multiple boundary repulsions within-subjects, for both risky decision making and perception. Context independent models like Prospect Theory^4,5^, stochastic expected utility theory^23^ or noisy outcome sampling models^43,44^ cannot account for the flexible induction of new weighting patterns. Context-dependent models that rely on long term assumptions for statistical distributions over probabilities^10,40^ also cannot account for sudden changes in distortions with simple instructions of bounded ranges to participants, as in our experiments. Existing Bayesian models^14,17,19^ and approximate Bayesian sampling models^49^ can account for these patterns in two possible ways. One is by assuming that while participants usually use priors with peaks centered at 0.5 (creating the characteristic probability weighting pattern), they can flexibly and immediately adopt priors with peaks centered in the relevant ranges we induce in our experiment. Existing Bayesian accounts^14,17,19^ use this assumption often in combination with log-odds encoding to account for probability weighting. The alternative, which we propose, is that bounded priors naturally reflect the limits of probabilities (0– 1) and adapt to newly imposed experimental ranges, thereby explaining both the classical weighting pattern and its systematic changes largely independent of specific encoding functions or prior shapes (see Methods, Supplementary Figs. 2–3). Moreover, the same qualitative features also emerge under bounded priors constraining efficient coding (see Methods, Supplementary Fig. 1). Thus, our cognitive boundary repulsion account provides a parsimonious mechanistic origin for probability distortions and their systematic modulation by newly induced boundaries.

### Inducing new boundaries reduces variability in lottery valuation and probability estimation close to the boundaries

A third prediction of our account is that behavioral variability should decrease systematically in proximity to boundaries, for both natural and experimentally induced boundaries (Fig. 6a, c, e). This signature arises from the truncation of the posterior due to truncation at the boundaries, leading to reduced variability in behavior (Fig. 1i). This effect should be stronger for higher levels of cognitive noise. Following our pre-registered analysis plan, we quantified variability as the average deviation in reported values or estimates from the mean at each given probability. Then we calculated a rolling average (window size = 0.1) of this variability measure across the probability range and compared this measure of variability within our pre-registered probability bins (bin size = 0.15; grey bins in Fig. 6) (see Methods for full details). We compared these measures of response variability across different blocks that only differed in the presence or absence of contextual boundaries, holding stimuli, noise levels, and response scales constant. This ensures that any changes in variability could arise only from changes in cognitive inference due to the presence of the boundaries.

**Fig. 6.**
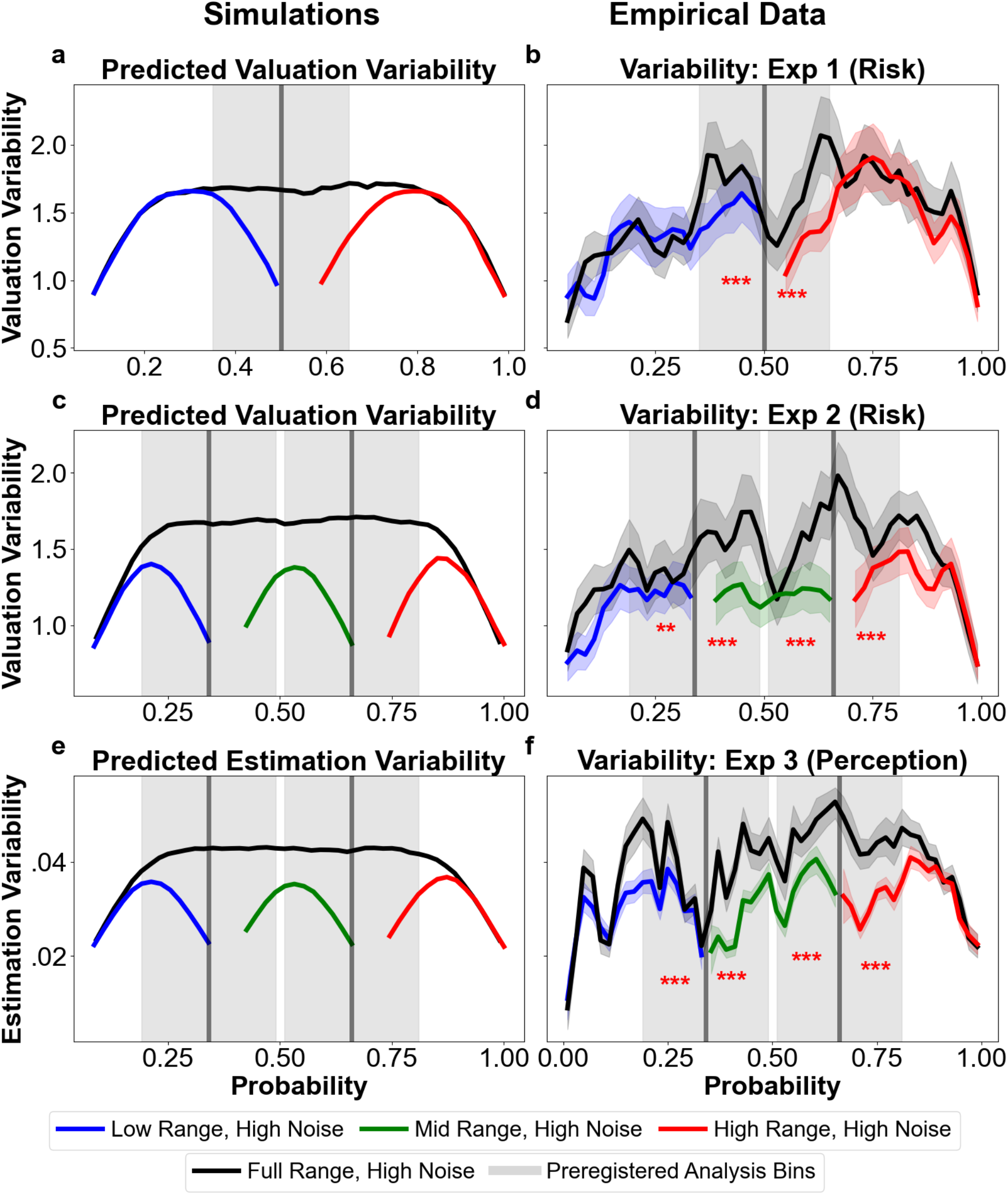
Effect of induced boundaries on variability in behavior: **a, c, e**. Model predictions of variability suppression due to cognitive boundary repulsions under high internal noise *(σ = 0*.*06)*. Simulations assume linear encoding and Bayesian decoding with a uniform bounded prior matched to the experimental block ranges (see Methods). For each condition, trialwise estimate variability was computed as the rolling standard deviation (window size = 0.1 in probability space) across the range. **a**. In Experiment 1 (lottery valuation), the model predicts reduced variability in certainty equivalents (CEs) near a contextual boundary at 0.5 in restricted-range blocks (red/blue) compared to the full-range block (black), due to posterior compression near the boundary. **c, e**. Simulated variability suppression in Experiments 2 and 3 when boundaries are added at 0.34 and 0.66. Model predicts local dips in variability around each induced boundary, holding noise constant. **b, d, f**. Empirical results confirm model predictions across all three experiments. **b**. In Experiment 1, introducing a contextual boundary at 0.5 under high noise significantly reduced variability in lottery valuations on both sides of the boundary. **d**. In Experiment 2, boundaries at 0.34 and 0.66 produced systematic reductions in variability around each boundary during valuation. **f**. In Experiment 3 (perceptual probability estimation), the same pattern was observed: variability in probability estimates decreased locally near both contextual boundaries. Grey shaded areas mark pre-registered probability bins used in analyses. Black lines show mean variability; grey bands show ±1 s.e.m.; red asterisks mark statistically significant effects in pre-registered bins (*p* < 0.05, \*\**p* < 0.001).

All three experiments confirmed our directional predictions of the effects of boundaries on variability in behavior, for both risky choice and simple perception. In *Experiment 1*, the introduction of a single contextual boundary at 0.5 led to a significant reduction in the variability of lottery valuations, on both sides of the introduced boundary (compared to blocks without this introduced boundary, both betas < 0, both p < 0.001; see Tables 4.1.1 and 4.2.1) (Fig. 6b). A dip in variability near 0.5 in the full-range condition most likely reflects rounding effects (i.e., participants tend to round off their estimates to values like 25, 50, or 75), but our within-subject design isolates the specific effect of boundary presence from such baseline effects, since they should occur in all conditions. In *Experiment 2*, adding two boundaries at 0.34 and 0.66 produced the corresponding reductions in variability on both sides of each boundary (all betas < 0, three p < 0.001, one p < 0.01; see supplementary Tables 4.1.2, 4.1.3, 4.2.2, 4.2.3) (Fig. 6d). These effects of reduced variability also generalized to perceptual judgments of probability in Experiment 3. Participants also showed significantly reduced variability near contextual boundaries (all betas < 0, all p < 0.001; see supplementary tables 4.1.4, 4.1.5, 4.2.4, 4.2.5) (Fig. 6f). Thus, the results of all experiments provide convergent evidence for the co-occurring signatures of distortions and variance patterns of cognitive boundary repulsions.

Our findings confirm that variability patterns, like average distortions, are systematically shaped by contextual boundaries—a finding that existing models cannot explain. Context-independent models either do not predict variability^4,5,42^ or would not predict changes in variability patterns due to these experimentally-induced boundaries^23,43,44^. Context-dependent models that rely on long term statistical distributions (prior shapes) over probabilities also cannot predict such rapid within-subject changes in variability patterns with simple boundary instructions to participants^10,40^. While the distortion patterns we get may in principle also be captured by a rapid change of priors to be of highest density in the center of the ranges, the localized variance reductions near induced boundaries can generally not be accounted for by such dynamic priors. Therefore, existing Bayesian models with flexible centrally shaped priors and fixed log odds encoding^17,19^ cannot account for these boundary induced variance changes. In contrast, our account can explain both biases and variance patterns, as well as their shift with induced boundaries, via the same unifying mechanism of cognitive boundary repulsions. This mechanism may be embedded in either Bayesian decoding with a bounded prior (Fig. 6) or in efficient encoding adaptation to bounded resources (Fig. 7d), which both result in changes in bias and variability. Thus, our boundary repulsion account provides a parsimonious mechanistic explanation for the joint contextual modulation of both distortions and variability across risky choice and perceptual probability estimation.

**Fig. 7.**
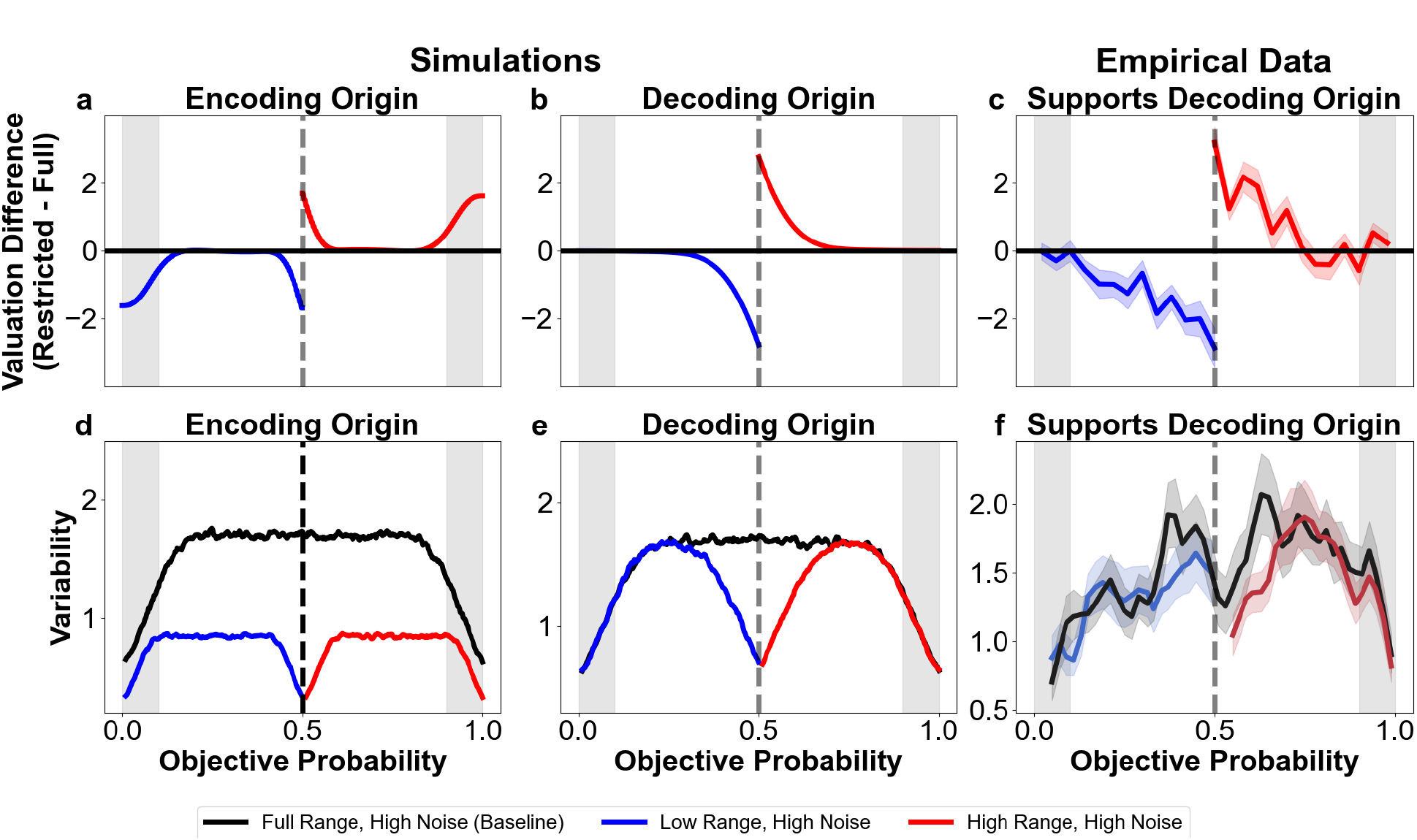
Origins of cognitive boundary repulsions at induced contextual boundaries: **(a)** When repulsions arise from efficient encoding, introducing a contextual boundary compresses the input range, causing encoding resources to adapt. This leads to a reduction in noise-driven distortions even at the original natural boundaries (shaded grey region), where biases due to boundary repulsions would otherwise be large. **(b)** In contrast, if repulsions emerge from Bayesian decoding with stable encoding, only the region near the new boundary is affected, and distortions in the grey region remain unchanged, showing range-independent effects. This divergence provides a key test: if encoding adapts, distortions at the original boundaries should shrink; if it doesn’t, they should stay the same. **(c)** Empirical data from Experiment 1 show that distortions near the original boundaries do not change with range manipulation— matching the prediction of decoding-based origins and providing no evidence for adaptive encoding in these regions. **(d)** Under efficient encoding, compressing the input range also leads to an overall decrease in variability, including in regions far from the newly introduced boundary (again, shaded grey). **(e)** In contrast, Bayesian decoding predicts that variability will dip only near the new contextual boundary, while remaining stable elsewhere, consistent with a non-adaptive encoding process. **(f)** Empirical variability data also show no range-dependent changes near the original boundaries, replicating the decoding-based signature and providing evidence against dynamic reallocation of representational resources (i.e, adaptive efficient coding).

### Do shifts in cognitive boundary repulsions originate at encoding or decoding?

Our manipulation of contextual boundaries elicited clear boundary repulsion effects: participants systematically overweighted probabilities above the newly introduced boundaries and underweighted those below, accompanied by a local reduction in variance near those boundaries. To identify whether these effects originated from adaptation at the encoding stage—via adapted efficient coding of the bounded input within the new range—or from Bayesian decoding using bounded priors, we leveraged the fact that both models make qualitatively different predictions about the effect of experimentally induced boundaries on response variability (see Methods).

While both mechanisms predict similar effects on variability near the new boundaries, they diverge in their predictions far away from the newly induced contextual boundaries. Under efficient encoding, the limited representational resources adapt to the smaller bounded range of inputs. As a result, the same level of internal noise in this encoded space now corresponds to a smaller range of inputs, and therefore, any noise-driven distortions and variability should scale with the bounded input range. A simple test of this prediction is to check the distortions and variability in behavior close to the natural boundaries of 0 and 1, far away from the new induced boundaries. If the noise-driven distortions and variability patterns that were present here due to the natural boundaries diminish for smaller ranges, this implies adaptive encoding and thereby an encoding-based origin for the newly introduced contextual boundaries. In Fig. 7a, the *y*-axis shows the simulated valuation difference (*restricted-range minus full-range*) under adaptive efficient coding with a uniform bounded prior and high noise (σ = 0.06). The negative difference in the grey regions means that the positive biases normally observed near 0 and 1 are smaller under restricted ranges than under full ranges. In contrast, with Bayesian decoding for uniform bounded priors and high noise (σ = 0.06) under non-adaptive encoding, distortions at the natural boundaries remain unchanged (Fig. 7b). Empirical data from Experiment 1 (Fig. 7c) match this decoding-based signature. The predictions for variability also differ. Under efficient encoding, compressing the range reduces variability globally, including near the natural boundaries (Fig. 7d). By contrast, Bayesian decoding predicts variability decreases only locally around the new boundaries, with no change in the grey regions (Fig. 7e). Empirical data (Fig. 7f) again support the decoding-based account: variability reductions occur only near induced boundaries, with no evidence for adaptive rescaling at the natural boundaries. Thus, the key empirical test is whether noise driven effects at the original boundaries are range-sensitive (encoding) or range-invariant (decoding).

To formally assess this, we conducted model comparisons using Bayes factors, evaluating whether including range changes due to contextually introduced boundaries (probRange) improved model fits for bias and variability in bins far away from the added boundary. This was done for all three experiments. Simulations and data from Experiment 1 are shown in Fig. 7 for illustration.

The results reveal that in almost all bins and experiments, Bayes Factors provided evidence that contextual range does not influence either bias or variability far from the new boundary. Across the 12 tests, *10 Bayes factors were < 0*.*3* (see Supplementary Table 5), supporting the null hypothesis of no contextual range effect. These findings offer converging evidence from both bias and variability patterns, supporting a decoding-based origin for the contextual boundary repulsions observed in our tasks. The only notable exception occurred in Experiment 2 (Risk Double) near 0, where both bias and variability exhibited robust evidence for range-dependent changes (BF_10_ >> 1; see Supplementary Table 5)—suggesting possible adaptation at the encoding stage in that specific condition. However, the absence of similar effects elsewhere suggests that cognitive boundary repulsions due to newly induced boundaries in our experiments were largely decoding-based in origin.

## Discussion

Why do people consistently overweight small probabilities and underweight large ones? Traditionally, this pattern has been understood as a characteristic of risky decision-making and has been firmly embedded in the dominant model of risky choice^4,5^. However, similar patterns have been observed in domains that involve no risk at all, including expected value judgments^50^ and in simple perceptual judgments of percentages^17,51^, and the cognitive origins and determinants of probability weighting are largely unclear. Here we propose that probability weighting originates from the cognitive processes involved in the inference of any naturally bounded variable, as often formalized within a two-stage encoding-decoding framework^9^. In contrast to other approaches, we do not propose that probability weighting reflects specific encoding functions or priors employed during this process, but instead arises from general and optimal computational processes that are foundational for the two stages: either mutual-information-maximizing encoding of bounded quantities or statistically optimal Bayesian decoding with bounded priors. We show that when the inferred quantities are bounded—such as probabilities between 0 and 1—both mechanisms can induce systematic repulsions in noise away from the boundaries. This leads to co-occurring predictions of the classic probability weighting pattern and a distinctive reduction in behavioral variability near boundaries. Our account of probability weighting is fully constrained, without requiring any fitted priors or functional assumptions to explain the major patterns, and instead relies solely on the known bounded structure of probabilities along with the cognitive noise that is inevitable in the brain’s information processing.

Our account predicts that both the shape and strength of probability weighting can be systematically modified by altering the boundaries and cognitive noise. It also predicts that similar patterns should occur across different domains of cognition, such as perceptual estimation and risky valuation tasks. Specifically, increasing cognitive noise should amplify the weighting pattern, while introducing new contextual boundaries should create additional weighting near the new boundaries. Crucially, the model predicts not only changes in bias but also distinctive dips in variability near both natural and induced boundaries, as diagnostic signature of cognitive boundary repulsions. We tested these predictions in three pre-registered, within-subjects experiments that manipulated two core factors: cognitive noise, by varying the complexity of numerical probability formats (e.g., simple vs. complex fractions), and boundary structure, by explicitly altering the range of probabilities presented within each block (e.g., 0–1, 0–0.5, 0.34– 0.66). Across all conditions, we observed results consistent with our predictions: both the strength and shape of probability weighting—and the associated variability—shifted systematically in response to our manipulations. These findings validate the core mechanism of boundary repulsion under noisy inference and demonstrate its generality across tasks.

Our work builds on a growing body of research that frames probability weighting not as a fixed, descriptive weighting function, but as an emergent consequence of noisy cognitive inference^10,14,17,19^. These models shift away from traditional descriptive models which treat the weighting curve as a static, parametric transformation of probability^5,42^. By modeling internal noise in encoding or transformation, recent approaches can explain how the strength of distortion may vary with cognitive load or task demands. However, they still rely on assumptions such as fixed log odds encoding^17,19^, encoding prior shapes^10^ or centrally anchored priors^14,17,19^ to reproduce the characteristic patterns. None of these assumptions can account for the large, boundary-sensitive shifts in valuation and perception (up to 15%) we observed when experimentally introducing novel stimulus boundaries. Moreover, these assumptions also cannot account for the dips in variability that we induced within-subjects with the boundary manipulations. Alternative explanations, including truncation of errors at the highest and lowest lottery outcomes^23^ and outcome-sampling models that assume noisy memory draws^43,44^, also cannot account for these empirical findings. Thus, probability weighting is best understood as a natural consequence of cognitive boundary repulsion—a parsimonious account that requires no additional assumptions, explains established patterns of distortions and variance, and uniquely predicts the novel boundary-sensitive phenomena we validate across three preregistered experiments.

Beyond risky choice and simple perceptual judgment, our account may be extended to offer a fresh lens on several other puzzling findings in economic behavior. For instance, the stake size effect—where individuals exhibit risk-seeking for small stakes and risk aversion for large ones^19,52^—may reflect boundary repulsions of values/utilities within the constrained range of experimental rewards (or biases due to a poorly learnt central prior). Similarly, findings on loss aversion show that changing the range (contextual boundaries) of gains and losses can alter behavior^53^, consistent with our claim that contextual boundaries lead to systematic biases in perception of key decision variables that propagate into higher order cognition. Further studies should test the applicability of our account to these empirical patterns. If confirmed, this would provide converging evidence for the emerging view that revealed preferences as measured with choice paradigms may partly reflect structured cognitive biases and not (just) intrinsic economic preferences^45^. Recognizing this distinction may help to resolve concerns about the instability of preferences and may contribute to more mechanistically accurate models of decision-making^54,55^.

Our work situates probability weighting within a broader class of perceptual and cognitive biases that emerge from noisy inference over bounded quantities. In perceptual neuroscience, central tendency effects in estimation and contraction biases in comparative judgments have been widely observed across domains such as length, numerosity, time, brightness, and even value^56–62^. These paradigms typically involve estimation of stimuli drawn from a constrained range, possibly forming contextual boundaries that - although not always explicitly instructed - are often implicitly learned through repeated exposure. The resulting biases, characterized by overestimation of small magnitudes and underestimation of large ones, closely mirror the boundary repulsion effects we observe and, like them, scale with cognitive noise. Traditionally, such biases have been attributed to Bayesian decoding with a central hump-shaped prior, which pulls estimates toward the mean of the stimulus distribution, thereby creating such central tendency effects^59^. However, other work in these perceptual domains has suggested that similar effects may also arise from decoding with uniform bounded priors, which create repulsions away from the boundaries^24–26^. Our account builds on this foundation, but extends it in two ways: first, by showing that boundary repulsions can emerge from either decoding or encoding processes in dissociable ways; and second, by explicitly manipulating boundary structure through instructions and demonstrating their causal role in shaping both probability weighting and variability in perception and valuation.

The distinction between central tendency and boundary-based biases raises a deeper question about how structure shapes inference: do people estimate stimulus values primarily from expectations about their generative distribution, or from knowledge of their boundaries? A recent study^26^ embedded both processes in a single computational framework, offering an important tool for adjudicating between these alternatives. We leverage this insight in our work: Using explicit boundary manipulations and testing both distortion and variability, we find patterns that are uniquely consistent with Bayesian decoding using bounded rather than central priors. Moreover, we extend this approach to a second level of dissociation, identifying not only what kind of prior is used in decoding, but also whether the observed biases arise from decoding or from resource-rational encoding. Our account predicts that encoding-based repulsions should scale with stimulus range, producing range-dependent biases and variability, whereas decoding-based repulsions should remain range-independent. This is generally consistent with prior work showing that behavior and neural representations adapt to stimulus ranges, which can be understood as a form of efficient encoding^31–39,47^. Our account builds on this view, by mapping this range-dependent signature back to the cognitive boundary repulsions that may arise due to adaptive efficient coding. However, across three experiments in two cognitive domains, our data reveal distortions and variability patterns that were largely range-independent (i.e., repulsions near the natural boundaries of 0 and 1 in the full range condition were indistinguishable from those observed when smaller ranges were induced), thereby supporting a decoding-based origin for boundary repulsions. Even partial encoding adaptation would have produced measurable range dependence, which we did not observe (except for one condition in Experiment 2). Thus, under the conditions we studied (probabilities expressed as complex fractions with explicitly instructed boundaries), encoding appeared to remain largely fixed and decoding appeared far more likely to explain the observed biases. Importantly, our account now offers a way to dissociate the origin of similar effects in other settings where encoding may adapt and produce range-dependent cognitive boundary repulsions, such as choices in different modalities, with longer training, or with implicitly learned boundaries. Future work should systematically vary these factors to clarify the timescales and task demands that may shape adaptive encoding in the brain.

Our findings also offer practical implications for both preference elicitation as well as policy intervention. With regards to preference elicitation, when the aim is to recover true preferences from observed choices, it is essential to account for distortions introduced by contextual boundaries. We show that boundary manipulations can shift valuations by more than 15% within individuals, without changing any objective feature of the decision. This highlights the importance of recognizing how elicitation methods can systematically influence measured preferences^63^. Once the cognitive mechanisms driving these distortions are well-understood, such biases become predictable and can be accounted for. For policy interventions, the same mechanistic understanding can be used constructively. Prior work shows that changing choice architecture or framing can robustly impact real-world choice behavior^64,65^. Our account identifies two concrete targets for policy interventions—reducing cognitive noise and reshaping perceived boundaries— that can stabilize decisions and align them more closely with people’s underlying goals and preferences.

In sum, our work provides a unifying explanation for the emergence of probability distortions in perception and risky choice, embedded in a single, mechanistic account of bounded noisy inference. In contrast to the prevailing view of probability weighting under risk as an irrational idiosyncrasy, we point out that the characteristic pattern of probability weighting emerges from optimal, resource-rational cognitive inference of bounded variables under cognitive noise. Critically, this account offers a tractable way to predict, measure, and intervene on cognitive biases across domains, from low-level perception to high-level economic decision-making. Looking ahead, this perspective opens the door to a richer understanding of how cognitive limits shape behavior and to new possibilities for bridging theories of perception, cognition, and economic choice.

## Methods

### Participants

In total, two-hundred-thirty-four (234) healthy young volunteers participated in the three experiments. Experiments were conducted on different days with non-overlapping groups of participants. Two pre-registered exclusion criteria were applied. First, for each subject, we combined data from all blocks and fit a linear regression of certainty equivalents (estimated probabilities) on presented probabilities. Data points more than 3 z-scores from the mean were excluded as outliers. This step aimed to remove values likely due to excessive mis-clicks or lapses of attention. Second, we fit separate linear regressions of certainty equivalents/estimated probabilities against probabilities for each run. Participants were excluded if any run yielded a non-positive slope or a slope not significantly greater than zero, as this indicates failure to integrate probability information meaningfully, suggesting random responding. Before exclusions, our sample sizes for the three experiments were 84 for experiment 1, 87 for experiment 2 and 63 for experiment 3. After applying the exclusion criteria, we reached the final sample comprising 200 participants consistent with our pre-registered target: 71 (age 19-32 years, 31 females) in Experiment 1 (risky valuation with one added boundary), 70 (age 19-37 years, 33 females) in Experiment 2 (risky valuation with two added boundaries), and 59 (age 19-33 years, 32 females) in Experiment 3 (perceptual probability estimation with two added boundaries). All participants provided written informed consent prior to participation. Participants received a base show-up fee of 10 CHF, plus additional monetary compensation between 0 and 40 CHF based on their choices in the risky valuation experiments or their accuracy in the perceptual estimation experiment. Testing sessions were conducted at the behavioral lab between 09:00 and 17:00. The study conformed to the Declaration of Helsinki and was approved by the Human Subjects Committee at the Faculty of Economics, Business Administration, and Information Technology at the University of Zurich.

### Procedure

All participants were tested in a behavioral lab equipped with multiple computers, with each participant seated at a separate workstation. Sessions began with informed consent, followed by written task instructions and a general set of comprehension questions to assess understanding of the task. Each participant completed one of three experiments: risky lottery valuation with one added boundary (Experiment 1), risky lottery valuation with two added boundaries (Experiment 2), or the perceptual probability estimation task (Experiment 3). In all three experiments there was also a control condition without any boundaries (except the natural ones at 0 and 1) – the full range. All tasks were block-structured, with each block defined by a specific probability range and noise level condition. Before each block, participants were explicitly informed of the relevant probability range and completed short comprehension checks to confirm understanding. Participants who answered comprehension questions incorrectly reviewed their responses with the experimenter and repeated the questions until all answers were correct. This procedure was enforced by an on-screen instruction requiring participants to call the experimenter by raising their hand; the screen could only be dismissed using a special key known to the experimenter. This step was critical, as the manipulation of cognitive boundaries in our experiment relied on participants being aware of the probability range presented in each block.

Across all experiments, probabilities were sampled uniformly within the instructed range in steps of 0.02 in probability space, starting at 0.01 and continuing up to the range’s upper bound. For example, full-range blocks covered 0.01 to 0.99 (49 values), lower and upper half-range blocks spanned 0.01–0.49 and 0.51–0.99 (25 values each), and narrower ranges used in Experiment 2 included 0.01–0.33 and 0.67–0.99 (17 values each), as well as a mid-range block from 0.35 to 0.65 (16 values). The number of repetitions per probability value and resulting trial counts per block differed between the lottery valuation and estimation tasks and are detailed in the corresponding sections below. Probabilities were presented in randomized order within each block. Block order was determined using a pre-registered sequential randomization procedure: participants were first randomly assigned to complete either the full-range or restricted-range portion of the task. Within the restricted-range portion, the order of low, high, and mid-range blocks (as applicable) was randomized. Within each range section, the order of the noise conditions— low versus high—were also independently randomized. In low-noise conditions, probabilities were presented as simple fractions over 100 (e.g., 32/100). In high-noise conditions, probabilities appeared as complex fractions composed of randomly generated four-digit numerators and denominators (e.g., 3724/8126). These complex fractions were generated prior to the experiment and were held constant across all participants and experiments. Fraction formats were introduced in the instructions and remained consistent within blocks.

Each session lasted approximately 1 hour and 15 minutes. In the risky lottery valuation tasks, participants were compensated based on one randomly selected trial per block, using a Becker– DeGroot–Marschak (BDM) auction procedure^66^. In the perceptual estimation task, payment was based on accuracy of probability judgments, as described below. All tasks were implemented using MATLAB R2024a with the Psychophysics Toolbox (Psychtoolbox) and presented on 24.5-inch monitors with a 1920 × 1080 pixel resolution and a 60 Hz refresh rate. All participants completed the tasks on identical setups in a controlled behavioral testing environment.

### Risky lottery valuation task

Experiments 1 and 2 employed a risky lottery valuation task in which participants indicated the minimum amount of money (certainty equivalent, CE) they would accept instead of playing a lottery offering a fixed payoff of 50 experimental currency units (ECU) with varying probabilities (Fig. 2a). Each trial began with a fixation cross displayed for 0.5 seconds, followed by the presentation of the lottery and response screen. Lotteries were presented on a standard computer monitor against a gray background, centered horizontally and shifted upward by 5% of the screen height, while participants’ certainty equivalents appeared at the exact screen center as they were entered. Participants had up to 9 seconds to enter their CE using number keys on the computer keyboard, for a total response window of 9.5 seconds. Invalid responses outside the 0–50 ECU range were flagged in red to allow correction. Once a response was submitted or the time expired, after a 0.5-second break, the next trial began. Both experiments used a two-factorial within-subjects design, crossing probability range and cognitive noise level (Fig. 2b–c). In *Experiment 1*, participants completed six blocks, crossing three probability ranges—full (0–1), low (0–0.5), and high (0.5–1)—with two noise levels: low noise (simple fractions over 100) and high noise (complex four-digit fractions). In *Experiment 2*, participants completed eight blocks with four probability ranges: full (0–1), lower (0–0.33), middle (0.35–0.65), and upper (0.67–1), crossed with the same two noise levels. In each block, every probability value sampled from the specified range was presented twice, resulting in approximately 100 trials per full-range block, 50 trials for half-range blocks, 34 trials for the narrow low and high blocks, and 32 trials for the mid-range block. Participants were allowed to enter any CE between 0 and 50 ECU, regardless of the probability range, to avoid mechanical boundary effects. At the end of the task, one trial was randomly selected from each block for payment. A *BDM auction procedure* was employed to determine the payout: the experiment computer generated a random offer between 0 and 50 ECU. If the participant’s CE was higher than the offer, they played the lottery; if same or lower, they received the computer’s offer instead. The average payoff across selected trials determined the participant’s final bonus.

### Perceptual estimation task

Experiment 3 used a perceptual estimation task in which participants judged visually presented probabilities and reported their estimates as percentages between 1 and 99% (Fig. 2d). Each trial began with a 0.5-second fixation cross, followed by the presentation of a fraction representing a probability. Fractions were presented on a standard computer monitor against a gray background, centered horizontally and shifted upward by 10% of the screen height, while participants’ probability estimates appeared at the exact screen center as they were entered. Participants had up to 5 seconds to type their estimate using number and decimal keys and confirmed their response by pressing Enter. Invalid responses outside the 1–99 range were flagged in red, allowing correction. Once submitted or timed out, the fixation cross briefly turned green, and after a 0.5-second break, the next trial began. The task followed a two-factorial within-subjects design, crossing probability range and noise level (Fig. 2e–f). Participants completed eight blocks comprising four probability ranges: full (0–1), lower (0–0.34), middle (0.34–0.66), and upper (0.66–1), crossed with two noise conditions: low (simple fractions over 100) and high (complex four-digit fractions). As in the valuation experiments, probabilities were uniformly sampled in steps of 0.02 and presented in random order. In *low-noise blocks*, each probability was presented once. In *high-noise blocks*, each probability was presented five times, allowing for a more reliable measurement of estimation variability. In the low-noise condition, participants could effectively respond by reading and converting the numerator of the simple fraction (e.g., the correct response for 34/100 was 34). Participants were instructed to be as accurate as possible. Payment was based on the error of their responses: For each trial, the absolute deviation between the participant’s estimate and the true probability was computed. The total deviation across trials was penalized using the formula: *bonus = 40 CHF – (total deviation × 2*.*5 / 600)*. A base compensation of 10 CHF was guaranteed, and no participant received less than this amount.

### Theory: Cognitive boundary repulsions from encoding and decoding

In the following, we formalize how cognitive boundary repulsions - biases away from the boundaries of a bounded quantity, accompanied by reduced variability near boundaries - emerge generically in a two-stage inference framework comprising noisy encoding followed by Bayesian decoding. These repulsions can arise independently from two distinct mechanisms.

First, under *efficient encoding*, limited representational capacity combined with mutual information maximization induces asymmetric, truncated likelihoods near the boundaries. These lead to *range-dependent boundary* repulsions.

Second, under *Bayesian decoding*, even with unconstrained or inefficient encoding, applying a *bounded prior* truncates the posterior, producing boundary repulsions in noise that are *range-invariant*.

These subtly different predictions of these two accounts of boundary repulsion yield distinct behavioral signatures, which we exploit to test the origin of distortions in our experiments.

### Boundary repulsions at the encoding stage: Resource-rational efficient coding

We model the noisy encoding process as:

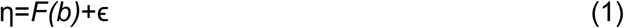

where *F*(*b*) is a deterministic encoding function mapping stimulus *b* ∈ [*b*_1_, *b*_2_] onto a bounded representational space η ∈ [0, *C*], ϵ is zero-mean, symmetric noise truncated to remain within the resource bounds [0, *C*].

#### Boundary repulsions due to efficient encoding

We assume that the encoding function *F*(*b)* is optimized to maximize mutual information *I*(*b*; *η*) between stimulus (*b)* and internal representation (η). Mutual information is:

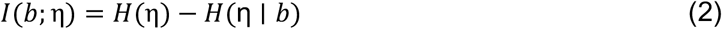

Because the representational space is bounded, noise is also truncated at 0 and *C*, introducing asymmetries in noise and the likelihood function due to truncation as soon as *F*(*b*) deviates from the center of the representational space (0.5C). At *F*(*b*) = 0.5*C*, truncation is symmetric, and so is the likelihood (*L*(*η* ∣ *b*)). At *F*(*b*) < 0.5*C*, negative noise is preferentially truncated, inducing positive skew in the likelihood. At *F*(*b*) > 0.5*C*, positive noise is preferentially truncated, inducing negative skew. These truncation-induced asymmetries emerge whenever the internal noise has sufficient spread relative to the bounds — as is typically the case with realistic noise distributions like Gaussian, Laplace, or Poisson. For such distributions, even small deviations of *F*(*b*) from the center of the representational space (i.e., 0.5C) lead to truncation-induced asymmetries in the likelihood. In contrast, for very narrow or strictly bounded noise (e.g., uniform over a small interval), these asymmetries only arise near the edges.

Due to truncation at [0, *C*], the likelihood function and conditional entropy satisfy mirror symmetry around the center point of the representational space (*F*(*b*) = 0.5*C)*:

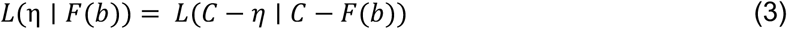

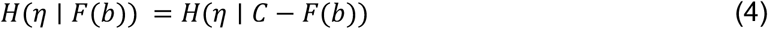

To formally show that mutual information maximization requires the encoding function *F(b)* to utilize both sides of the encoding space around 0.5*C*, assume for contradiction that the optimal encoding function only spans one half: *F*(*b*) ∈ [*x, y*] ⊂ [0*C*, 0.5*C*]. Then, define a mirrored encoding function F’(b) = C − F(b) which spans the corresponding region on the other half [C − *x*, C − *y*] ⊂ [0.5*C, C*]. By equation (3), the conditional entropy is the same for F and F’. However, combining both mirrored encodings F and F’ would cover a wider span of the representational space increasing the **marginal entropy** *H*(η). This contradicts our initial assumption of maximizing mutual information, given in equation (2). Therefore, the optimal encoding (*F*) must span both sides of the representational space around 0.5*C* to maximize mutual information. Since truncation-induced asymmetries on the two sides of 0.5*C* induce inward skew, this result implies a general consequence. When a bounded quantity is encoded into a limited capacity representational space under mutual information maximization, the resulting representations exhibit cognitive boundary repulsions. This leads to an overestimation for small quantities transitioning to underestimation for large ones, where the transition point depends on the shape of the prior distribution constraining the encoding function under efficient coding (see Supplementary Fig. 1 for numerical simulation examples showing this).

While our theoretical derivations define efficient coding as maximizing mutual information between stimuli and internal representations^13,46,67,68^, exact closed-form solutions are generally unavailable because MI optimization is analytically intractable for most realistic noise distributions. In practice, MI-optimal solutions are often approximated using Fisher information under additional assumptions about the noise structure^67,68^. For our simulations, we therefore implemented efficient encoding using the cumulative distribution function (CDF) transform of the prior, a classic redundancy-reducing code^12^ that equalizes responses and, under small symmetric noise assumptions, closely approximates MI-optimal encoding^13,67^. Even though with truncated Gaussian noise, the CDF transform is not strictly a mutual information maximizing efficient code, it nonetheless provides a principled and widely used approximation that captures the qualitative signatures of efficient coding. Our simulations with this approach robustly reproduced the theoretically predicted boundary repulsion effects, supporting our theoretical claims (Supplementary Fig. 1).

### Range dependence of cognitive boundary repulsions due to efficient coding

Let the original bounded stimulus be *b* ∈ [*b*_1_, *b*_2_] and the encoding function *F*(*b*): [*b*1, *b*2] → [0, *C*] be efficient in the sense of maximizing mutual information according to the prior distribution (*p*(*b*)). Now, consider linearly stretching the input range by a factor *k* > 0, such that *(b*′ = *k* · *b*), and *b*′ ∈ [*kb*_1_, *kb*_2_]. We assume that the prior scales accordingly, preserving its shape: 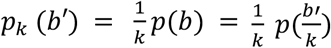.

Since the efficient encoder is defined based on the prior, the new encoding function over the stretched range satisfies 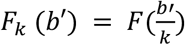. This simply says that encoding a stretched input b’ is equivalent to encoding the original unscaled value 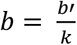.

Differentiating both sides with respect to b’ gives:

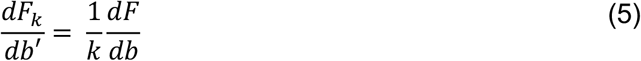

Equation 5 implies that the slope of the encoding function with respect to the stretched input b’ is scaled down by a factor of 1/k. That means the encoding becomes less sensitive (flatter) to inputs when *k* > 1 and more sensitive (steeper) when *k* < 1. Now consider how the same levels of cognitive noise in this new encoded space affect the corresponding perturbations in the stimulus space.

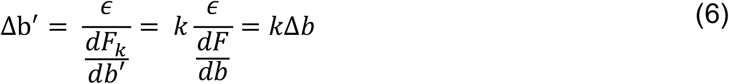

Hence, both bias and variance in external stimulus estimates scale linearly with the stimulus range under efficient encoding. Expanding the input range forces internal resources to stretch thinner, reducing local precision and amplifying both bias and variability in behavior for the same levels of cognitive noise.

### Conclusion

Efficient encoding of bounded quantities leads to inward-skewed likelihoods near the boundaries, producing cognitive boundary repulsions. Crucially, these distortions scale with the external range of input: expanding the stimulus range increases both the magnitude of bias and variability. This result holds generically for any prior distribution that stretches proportionally but conserves its shape with the stimulus range. Thus, the range dependence of boundary repulsions is a robust consequence of resource-rational efficient coding under bounded capacity.

### Cognitive boundary repulsion in posterior: Bayesian decoding

Even in the absence of efficient coding constraints, cognitive boundary repulsions can emerge naturally at the *decoding stage* when Bayesian decoding is applied to bounded quantities. Specifically, when internal representations η are decoded via Bayes’ rule using a prior p(b) defined over a bounded domain *b* ∈ [*b*_1_, *b*_2_], the resulting *posterior distribution is necessarily* truncated near the boundaries, inducing distortions in the decoded estimates. The encoding is still given by equation 1 defined above where *F*(*b*) is a deterministic, monotonic encoding function (here not necessarily efficient), and ϵ is zero-mean, symmetric noise (e.g., Gaussian, or any other zero-mean, symmetric noise distribution) that is independent of *b*. Given that limited representational resources are no longer assumed, the noise and likelihood are taken to be symmetric and untruncated. Now, at decoding, the brain computes the posterior via:

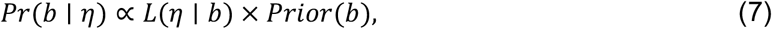

where the *Prior(b)* is bounded such that *Prior(b)*> 0 for *b* ∈ [*b*_1_, *b*_2_], and *Prior(b)* = 0 otherwise. Multiplying the likelihood by the bounded prior truncates the posterior at *b*_1_, and *b*_2_. Thus, the posterior distribution exhibits *inward boundary repulsions* even when encoding noise is untruncated and independent of *b*. This again leads to an overestimation for small quantities transitioning to underestimation for large ones where the transition point from right to left skew depends on the shape of the prior distribution and the encoding function (see Supplementary Figs. 2, 3).

### Range invariance of decoding-based boundary repulsions

As shown above, even when encoding is non-adaptive and does not impose resource-based truncations in the likelihood, Bayesian decoding of bounded quantities still leads to cognitive boundary repulsions through posterior truncation. Crucially, unlike encoding-based cognitive boundary repulsions, the boundary repulsions arising from Bayesian decoding do not scale with the stimulus range. When the entire input space is linearly stretched by a factor k, and encoding does not efficiently adapt to this new input range, the likelihood remains unchanged. This implies that the same levels of encoding noise led to the same corresponding perturbations in the stimulus space. The bounded prior only plays a role during decoding to truncate the posterior, inducing cognitive boundary repulsions. However, since the likelihood is unchanged and the posterior still maps to the same corresponding input, the corresponding perturbations in behavior remain range-invariant. Therefore, while both biases and variance are range-dependent for encoding based cognitive boundary repulsions, they remain range invariant when they arise entirely during decoding. This distinction offers a critical theoretical and empirical dissociation between the two origins.

## Simulations

All model simulations in Figs. 3–6 are based on Bayesian decoding of noisy internal representations derived from linearly encoded, bounded input probabilities with symmetric, untruncated noise during encoding. Bayesian decoding was applied using a uniform prior truncated to the contextual range of probabilities for each condition (e.g., [0–0.5], [0.5–1], [0.34–0.66]), reflecting the actual empirical prior distribution. Posterior means were computed for each internal representation to obtain predicted estimates. Two fixed levels of internal noise were used across all simulated predictions across all datasets: a low-noise condition (σ=0.02) and a high-noise condition (σ=0.06). These values were not fit to individual datasets or varied across simulations; they were chosen to match empirical behavioral variability and held constant across all experiments and domains. In valuation simulations (Experiments 1 & 2), inferred probabilities were simply multiplied by a fixed reward magnitude (v = 50 ECU) to generate predicted certainty equivalents. *Variability plots* (e.g., Fig. 6) were generated by taking two posterior samples for each probability level and computing the absolute deviation of those samples from the mean of the sampled estimates for that probability 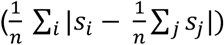. This process was repeated over a large number of simulated subjects (N = 1000 repetitions), and the mean deviation across simulated participants was computed for each probability. These deviations were then smoothed using a rolling average (window size = 0.1 in probability space) to produce the final predicted variability profile for each condition. This method closely mirrors the empirical calculation used in the data analyses, where observed deviations were computed relative to the empirical mean estimate at each probability.

For fig. 7, comparing encoding- and decoding-based origins of boundary repulsions under contextual range manipulations, parameter values used were identical to those used for simulations in Figs. 3–6. For the decoding-origin simulations, probabilities were linearly encoded with additive Gaussian noise and decoded with a uniform prior truncated to the relevant range ([0–1], [0–0.5], [0.5–1]). For the encoding-origin simulations, efficient coding was approximated by the normalized CDF of the uniform prior within the relevant range. Noise was modeled as Gaussian truncated to the representational bounds and decoding again used a uniform prior over the contextual range. Bias was computed as the difference in posterior means between restricted- and full-range conditions (*restricted-range minus full-range*) (Fig. 7a–b), and variability as mean of absolute deviations of posterior samples from the sample mean 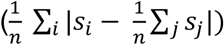 (N = 1000 repetitions; Fig. 7d–e). Encoding-origin simulations predict range-dependent changes at the natural boundaries, while decoding-origin simulations predict range-invariant effects, matching our theoretical predictions.

## Data analysis

All behavioral analyses were conducted using pre-registered generalized linear mixed-effects models of the form:

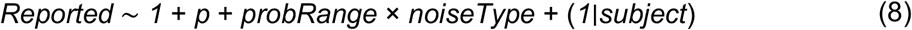

where *Reported* denotes either the certainty equivalent (valuation tasks) or the probability estimate (estimation task), depending on the experiment. This model tested the effects of objective probability (*p*), contextual boundaries (*probRange*), cognitive noise (*noiseType*), and their interaction, with random intercepts for subjects to capture individual variability.

To avoid overfitting trial-level noise and to focus on theoretically relevant regions, all tests were performed within pre-registered probability bins. For distortions (Hypotheses 1–3), bins of width 0.1 in probability space were defined adjacent to natural and contextual boundaries (e.g., [0.0– 0.1], [0.9–1.0], [0.4–0.5], [0.5–0.6]). For *Hypothesis 4*, which tested predictions about *behavioral variability*, the same model structure was used, but with the *rolling average of trial-wise absolute deviations* from the mean (variability) as the dependent variable. As the variability measure is noisier by nature, wider bins of width 0.15 were pre-registered (e.g., [0.35–0.5], [0.5–0.65]). Exact mixed-effects model results for all hypotheses are reported in Supplementary Tables 1–4. Our four pre-registered hypotheses were:

Hypothesis 1 was to test the effect of cognitive noise near the natural probability boundaries. The factor of interest was *noiseType*. We predicted that high cognitive noise would amplify distortions at the boundaries of 0 and 1. To test this, we examined the effect of *noiseType* within the bins [0.0–0.1] (above 0) and [0.9–1.0] (below 1). The results confirmed the prediction, showing significant overweighting above 0 and underweighting below 1 under high noise (see Supplementary Table 1 for exact model outputs).

Hypothesis 2 addressed the effect of contextual boundaries. Here we predicted that introducing new boundaries would alter distortions, leading to reduced estimates below a boundary and increased estimates above it. This was tested as the effect of *probRange* in bins adjacent to induced boundaries, for example [0.4–0.5] and [0.5–0.6] for a boundary at 0.5. The results showed strong and significant shifts in the expected directions, supporting the prediction (see Supplementary Table 3).

Hypothesis 3 focused on the interaction between contextual boundaries and cognitive noise. We predicted that distortions induced by contextual boundaries would be stronger when cognitive noise was high. This was tested as the *probRange × noiseType* interaction in bins adjacent to the induced boundaries. The results confirmed this prediction, showing robust interaction effects in the expected direction (see Supplementary Table 2).

Hypothesis 4 examined the effect of boundaries on variability. For this hypothesis, the dependent measure was trial-wise variability, modeled as

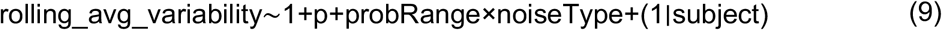

We predicted that variability would decrease near contextual boundaries under high noise. This was tested as the effect of *probRange* in preregistered bins of width 0.15 around induced boundaries. The results again supported the prediction, showing significant within-subject reductions in variability near all contextual boundaries (see Supplementary Table 4).

## Acknowledgements

We are grateful to Cornelia Schnyder and Elena Boldin at the Zurich Center for Neuroeconomics for their excellent assistance with recruitment and participant facilitation. C.C.R. received funding from the University Research Priority Program *Adaptive Brain Circuits in Development and Learning* (URPP AdaBD) at the University of Zurich, as well as from the Swiss National Science Foundation (SNSF, grant no. 100019L-173248). G.d.H. was supported by the Dutch Research Council (NWO, Rubicon grant no. 019.183SG.017/8O3B) and the University of Zurich (Forschungskredit grant no. K-33153-02-01). SB was supported by a PhD scholarship from the Marlene Porsche Graduate School of Neuroeconomics and funding by the URPP AdaBD. We also thank the organisers, participating faculty, and students of the 2023 Summer School of Neuroeconomics at the University of Pennsylvania, in particular Vaidyanathan Viswanathan and Camilla van Geen, for helpful discussions and feedback on an initial version of a pilot experiment testing these ideas.

