## Supplementary material for "Probability weighting arises from boundary repulsions of cognitive noise"

#### Supplementary note 1: Boundary repulsions arise during encoding across different prior shapes constraining efficient encoding

Supplementary Fig. 1 illustrates that the characteristic distortions in subjective probability inference (overestimation of small probabilities and underestimation of large probabilities) robustly emerge due to efficient encoding constrained by a wide range of bounded prior distributions. The simulations include U-shaped, hump-shaped, uniform, and skewed variants of these distributions (Supplementary Fig. 1a). Efficient coding is approximated as the cumulative distribution function (CDF) for each prior. This procedure adapts the representational mapping to the bounded probability space according to the prior distribution, and the resulting likelihood is then truncated at the resource bounds. In all simulations for this figure, decoding is performed using a uniform prior. All prior shapes lead to the same qualitative bias/distortion patterns (Supplementary Fig. 1b), demonstrating the robustness of the characteristic probability weighting pattern across different prior shapes.

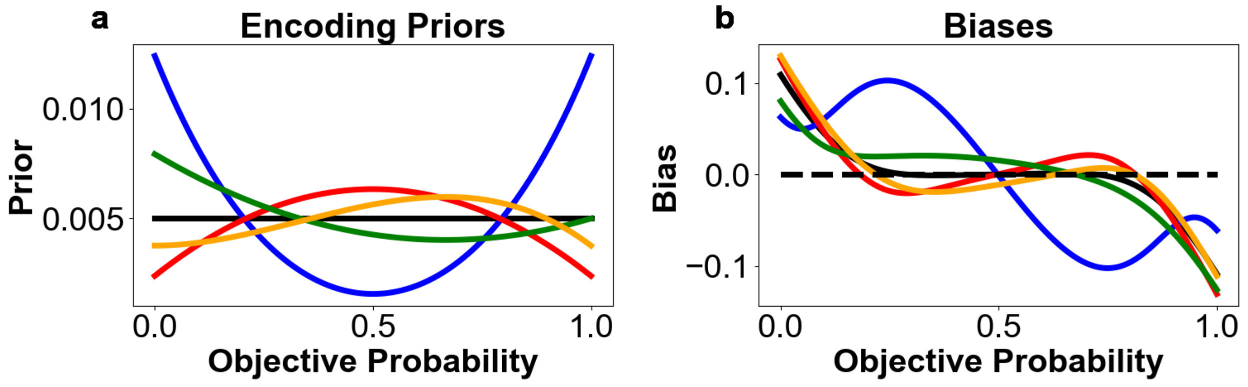


**Supplementary Fig. 1: Robustness of distortions due to efficient encoding under different encoding priors. (a)** Prior distributions used to constrain efficient encoding: uniform, U-shaped, hump-shaped, and skewed variants. Efficient encoding functions are produced by the cumulative distribution function (CDF) of each prior on the probability range [0, 1]. **(b)** Resulting biases in subjective probability inference under a uniform decoding prior. The y-axis shows bias = posterior mean estimate − objective probability, with additive truncated Gaussian noise (σ = 0.1) applied during encoding. Despite differences in the encoding prior, all priors yield systematic overestimation of small probabilities and underestimation of large probabilities demonstrating the robustness of the pattern to specific prior shapes. Thus, similar cognitive boundary repulsions emerge in general when bounded representational resources are efficiently adapted to bounded input quantity space during encoding.

### Supplementary Note 2: Boundary Repulsions Arise During Decoding Under Uniform Decoding for Different Encoding Functions

Supplementary Fig. 2 demonstrates that characteristic distortions in subjective probability estimation also emerge even when encoding does not adapt its limited representational space to the bounded space of quantities being encoded, due to Bayesian decoding (even with a uniform bounded prior). In these simulations, we show that a wide range of encoding function shapes—such as linear, concave, convex, or mappings that resemble log odds—all consistently give rise to the pattern of overestimation of small probabilities and underestimation of large probabilities. The exact curvature of these mean biases varies depending on the encoding function used, but the general distortion pattern due to boundaries remains robust. This highlights that boundary-induced distortions can emerge even when encoding is not optimized for information efficiency, but decoding is done with a prior representing only the bounded nature of the quantities being inferred.

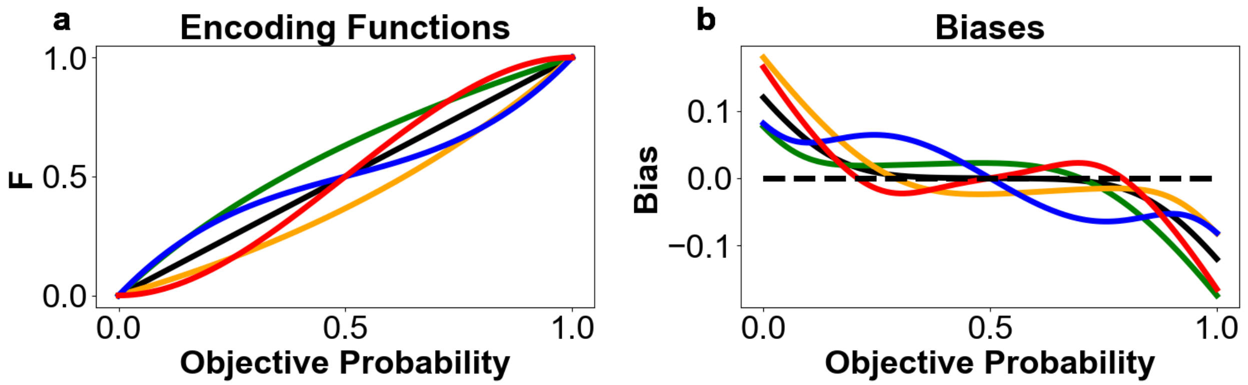


**Supplementary Fig. 2: Robustness of distortions due to Bayesian decoding for different encoding functions. (a)** Encoding functions tested: linear, concave, convex, and log-odds–like mappings of objective probability. Encoding was non-adaptive and included additive Gaussian noise (σ=0.1) without truncation. **(b)** Resulting **bias** in subjective probability inference under Bayesian decoding with a uniform prior bounded to [0, 1]. The *y*-axis shows *bias = posterior mean estimate − objective probability*. Despite differences in encoding curvature, all functions yield the characteristic pattern of overestimation at small probabilities and underestimation at large probabilities, demonstrating that boundary-induced distortions arise robustly from decoding with uniform bounded priors.

#### Supplementary Note 3: Boundary Repulsions Arise During Decoding Under Different Bounded Decoding Prior Shapes for Linear Encoding

Supplementary Fig. 3 demonstrates that the characteristic distortions in subjective probability estimation arise robustly across a wide range of bounded decoding prior shapes, even when the encoding function is fixed and linear. In these simulations, the representational mapping at the encoding stage does not adapt to the structure of the probability space, thereby not producing any boundary repulsions during encoding; instead, it linearly maps inputs to internal representations without any truncation. The variability in distortion patterns emerges solely from differences in the shape of the bounded prior used during Bayesian decoding. We simulate decoding under several bounded prior distributions, including uniform, U-shaped, hump-shaped, and skewed variants. While the exact curvature of biases differs depending on the prior's shape, all conditions consistently reproduce the classic pattern of overestimation of small probabilities and underestimation of large probabilities. This confirms that bounded decoding alone—even with a non-adaptive linear encoding—is sufficient to produce the boundary repulsion effects observed in human probability judgments, while the shape of the decoding prior primarily modulates the specific form and asymmetry of the distortions.

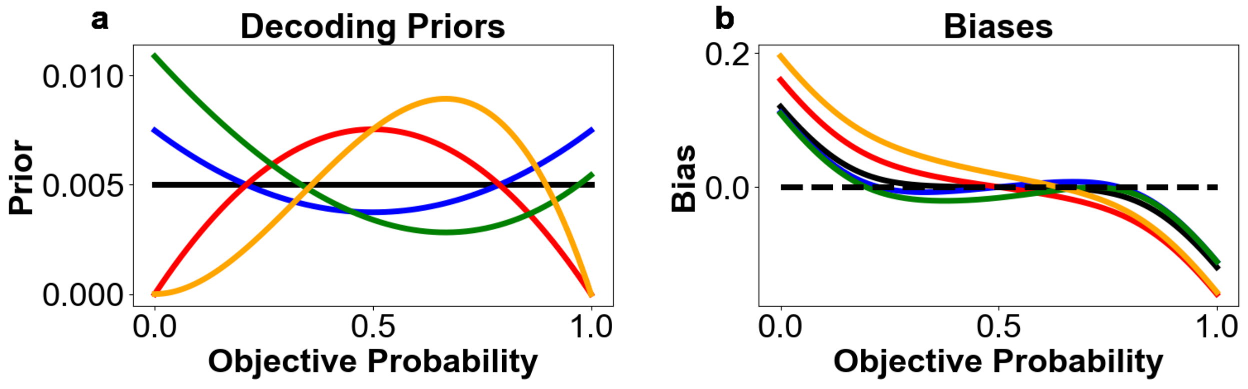


**Supplementary Fig. 3: Robustness of distortions due to Bayesian decoding under different bounded decoding priors for linear encoding. (a)** Prior distributions used for Bayesian decoding: uniform, U-shaped, hump-shaped, and skewed variants, all bounded on the probability range $[0, 1]$. **(b)** Resulting biases in subjective probability inference when inputs are linearly encoded (with additive Gaussian noise, σ = 0.1) and decoded with these bounded priors. The y-axis plots bias = posterior mean estimate − objective probability. Despite differences in the curvature of distortions across prior shapes, all decoding priors reproduce the same qualitative pattern of overestimation of small probabilities and underestimation of large probabilities. This demonstrates that cognitive boundary repulsions can arise purely from bounded Bayesian decoding, even when the encoding stage is fixed and non-adaptive.

### Supplementary Note 4: Pre-registered Mixed Effects Model Result Tables

#### Amplifying Probability Weighting by Increasing Cognitive Noise

The following tables show the results of analyses testing Prediction 1, that increasing cognitive noise amplifies the classic probability distortion pattern. As specified in our preregistration, we fitted linear mixed-effects models with participant as a random effect. Each model included fixed effects for the intercept, probability (p), noise type (low vs. high), probability range (depending on added boundaries in different blocks), and the interaction of noise and probability range. We focused on two preregistered probability bins: (1) probability bin between probabilities of 0 and 0.1 (above the lower natural boundary), and (2) the probability bin between 0.9 and 1 (below the upper natural boundary). We hypothesized that higher noise will lead to greater overestimation of small probabilities (reflected in positive beta regression values) and greater underestimation of high probabilities (reflected in negative beta regression values). Results for each experiment are reported in separate tables below. Experiment 1 and 2 involved risky lottery valuation and experiment 3 involved judging fractions denoting probabilities. Mixed-effects models are applied to either certainty equivalents (CE) or probability equivalents (PE), depending on the experiment.

##### Amplification of Overestimation Above the Lower Natural Boundary (0)

Tables 1.1.1, 1.1.2, and 1.1.3 show that increasing cognitive noise in probability inference leads to greater overestimation or overweighting of small probabilities (in our pre-registered bin of analysis). The key term of interest in these models is noiseTypeHighNoise, and the predicted effect is a positive regression coefficient, reflecting the hypothesized amplification.

#### Table 1.1.1: Experiment 1

Dependent variable: certaintyEquivalent. Analysis Bin: 0.00 to 0.10.
Reference levels: probRange = '00to100', noiseType = 'lowNoise'.

| term | estimate | std.error | t-statistic | df | p.value |
|---|---|---|---|---|---|
| (Intercept) | 0.940 | 0.256 | 3.674 | 107.1674 | 0.000187*** |
| p | 39.730 | 1.615 | 24.606 | 2,699.1875 | < 2e-16*** |
| probRange00to050 | 0.087 | 0.128 | 0.684 | 2,699.0090 | 0.246970 |
| **noiseTypehighNoise** | **1.600** | **0.129** | **12.438** | **2,699.1565** | **< 2e-16*** ** |
| probRange00to050:noiseType highNoise | -0.239 | 0.182 | -1.312 | 2,699.0604 | 0.094745. |

#### Table 1.1.2: Experiment 2

Dependent variable: certaintyEquivalent. Analysis Bin: 0.00 to 0.10.
Reference levels: probRange = '00to100', noiseType = 'lowNoise'.

| term | estimate | std.error | t-statistic | df | p.value |
|---|---|---|---|---|---|
| (Intercept) | 1.380 | 0.316 | 4.373 | 90.57995 | 1.63e-05*** |
| p | 43.823 | 1.628 | 26.925 | 2,649.82516 | < 2e-16*** |
| probRange00to033 | 0.022 | 0.129 | 0.166 | 2,649.83870 | 0.434 |
| **noiseTypehighNoise** | **2.642** | **0.130** | **20.250** | **2,649.97636** | **< 2e-16*** ** |
| probRange00to033:noiseType highNoise | -1.442 | 0.184 | -7.826 | 2,649.89952 | 3.62e-15*** |

#### Table 1.1.3: Experiment 3

Dependent variable: probabilityEquivalent. Analysis Bin: 0.00 to 0.10.
Reference levels: probRange = '00to100', noiseType = 'lowNoise'.

| term | estimate | std.error | t-statistic | df | p.value |
|---|---|---|---|---|---|
| (Intercept) | -0.081 | 0.508 | -0.160 | 207.764 | 0.437 |
| p | 102.370 | 3.386 | 30.232 | 3,211.689 | <2e-16*** |
| probRange00to033 | -0.037 | 0.454 | -0.082 | 3,211.141 | 0.467 |
| **noiseTypehighNoise** | **3.621** | **0.355** | **10.201** | **3,211.557** | **<2e-16*** ** |

Dependent variable: probabilityEquivalent. Analysis Bin: 0.00 to 0.10.
Reference levels: probRange = '00to100', noiseType = 'lowNoise'.

| term | estimate | std.error | t-statistic | df | p.value |
|---|---|---|---|---|---|
| probRange00to033:noiseType highNoise | -0.367 | 0.501 | -0.732 | 3,211.662 | 0.232 |

#### Amplification of Underestimation Below the Upper Natural Boundary (1)

Tables 1.2.1, 1.2.2, and 1.2.3 show that increasing cognitive noise in probability inference leads to greater underestimation or underweighting of large probabilities (in our pre-registered bin of analysis). The key term of interest in these models is noiseTypeHighNoise, and the predicted effect is a **negative regression coefficient**, reflecting the hypothesized amplification.

##### Table 1.2.1: Experiment 1

Dependent variable: certaintyEquivalent. Analysis Bin: 0.90 to 1.00.
Reference levels: probRange = '00to100', noiseType = 'lowNoise'.

| term | estimate | std.error | t-statistic | df | p.value |
|---|---|---|---|---|---|
| (Intercept) | -9.992 | 1.921 | -5.202 | 2,314.042 | 1.07e-07*** |
| p | 59.283 | 1.921 | 30.861 | 2,703.040 | < 2e-16*** |
| probRange50to100 | -0.132 | 0.154 | -0.857 | 2,703.080 | 0.196 |
| **noiseTypehighNoise** | **-2.472** | **0.154** | **-16.027** | **2,703.049** | **< 2e-16*** ** |
| probRange50to100:noiseType highNoise | 0.338 | 0.218 | 1.547 | 2,703.066 | 0.061. |

##### Table 1.2.2: Experiment 2

Dependent variable: certaintyEquivalent. Analysis Bin: 0.90 to 1.00.
Reference levels: probRange = '00to100', noiseType = 'lowNoise'.

| term | estimate | std.error | t-statistic | df | p.value |
|---|---|---|---|---|---|
| (Intercept) | -9.740 | 1.574 | -6.188 | 2,745.694 | 3.51e-10*** |
| p | 59.428 | 1.632 | 36.405 | 2,677.095 | < 2e-16*** |
| probRange66to100 | 0.048 | 0.131 | 0.364 | 2,677.088 | 0.358 |

Dependent variable: certaintyEquivalent. Analysis Bin: 0.90 to 1.00.
Reference levels: probRange = '00to100', noiseType = 'lowNoise'.

| term | estimate | std.error | t-statistic | df | p.value |
|---|---|---|---|---|---|
| **noiseTypehighNoise** | **-2.219** | **0.131** | **-16.912** | **2,677.098** | **< 2e-16*** ** |
| probRange66to100:noiseTypehighNoise | -0.009 | 0.185 | -0.047 | 2,677.109 | 0.481 |

Table 1.2.3: Experiment 3

Dependent variable: probabilityEquivalent. Analysis Bin: 0.90 to 1.00.
Reference levels: probRange = '00to100', noiseType = 'lowNoise'.

| term | estimate | std.error | t-statistic | df | p.value |
|---|---|---|---|---|---|
| (Intercept) | -18.935 | 3.157 | -5.998 | 3,412.912 | 1.11e-09*** |
| p | 119.931 | 3.285 | 36.513 | 3,356.609 | < 2e-16*** |
| probRange66to100 | 0.000 | 0.450 | 0.000 | 3,356.518 | 0.500 |
| **noiseTypehighNoise** | **-5.548** | **0.350** | **-15.837** | **3,356.707** | **< 2e-16*** ** |
| probRange66to100:noiseTypehighNoise | 0.545 | 0.495 | 1.100 | 3,356.614 | 0.136 |

#### Inducing New Boundaries Creates New Probability Distortion Patterns (Interaction Effects of Boundaries and Cognitive Noise)

The following tables show the results of analyses testing Prediction 2, which proposes that artificially introducing new boundaries in the probability space alters subjective probability distortions in predictable ways. In line with our preregistration, we fitted linear mixed-effects models with participant included as a random effect. Each model included fixed effects for the intercept, probability (p), noise type (low vs. high), probability range (reflecting contextually introduced boundaries across blocks), and the interaction between noise type and probability range. We focused on specific probability bins adjacent to these introduced boundaries, chosen based on preregistered criteria. These included regions just below and above $p = 0.5$ in Experiment 1, and just below and above $p = 0.34$ and $p = 0.66$ in Experiments 2 and 3. We predicted that cognitive noise would amplify the distortion in these regions for blocks that have boundaries, resulting in greater overestimation of probabilities just above, and greater underestimation just below these new bounds. This would be reflected in a positive regression coefficient above introduced boundaries and negative regression coefficient below due to the interaction between high noise and the contextual boundary manipulation. The results of these analyses are reported separately for each experiment. As in Prediction 1, Experiments 1 and 2 involved risky lottery valuation tasks and analyses were conducted on certainty equivalents (CE), while Experiment 3 involved judgments of explicit probability fractions and analyses were conducted on probability equivalents (PE).

#### Overestimation Above Contextually Induced Boundaries with Added Noise

Tables 2.1.1 through 2.1.5 show that introducing new contextual boundaries in the probability space leads to a systematic amplification of overestimation for probabilities just above those boundaries, under conditions of high cognitive noise when compared to when no boundaries are present there and low cognitive noise. As predicted, this effect is reflected in positive regression coefficients for the interaction term probRange:noiseTypeHighNoise. The observed amplification is consistent with the hypothesis that cognitive noise interacts with newly induced structure in the probability space to enhance distortions in subjective probability representation.

##### Table 2.1.1: Experiment 1

Dependent variable: certaintyEquivalent. Analysis Bin: 0.50 to 0.60.
Reference levels: probRange = '00to100', noiseType = 'lowNoise'.

| term | estimate | std.error | t-statistic | df | p.value |
|---|---|---|---|---|---|
| (Intercept) | 9.888 | 1.350 | 7.323 | 1,944.908 | 1.77e-13*** |
| p | 28.933 | 2.286 | 12.654 | 2,685.127 | < 2e-16*** |
| probRange50to100 | 0.231 | 0.179 | 1.286 | 2,685.102 | 0.0992. |
| noiseTypehighNoise | 0.048 | 0.180 | 0.268 | 2,685.107 | 0.3945 |
| **probRange50to100:noiseTypehighNoise** | **1.471** | **0.255** | **5.763** | **2,685.127** | **4.60e-09*** ** |

#### Table 2.1.2: Experiment 2 (First Added Boundary)

Dependent variable: certaintyEquivalent. Analysis Bin: 0.35 to 0.45.
Reference levels: probRange = '00to100', noiseType = 'lowNoise'.

| term | estimate | std.error | t-statistic | df | p.value |
|---|---|---|---|---|---|
| (Intercept) | 1.896 | 0.788 | 2.407 | 2,099.782 | 0.00809** |
| p | 45.441 | 1.875 | 24.237 | 2,902.233 | < 2e-16*** |
| probRange33to066 | 0.269 | 0.171 | 1.569 | 2,902.201 | 0.05835. |
| noiseTypehighNoise | -0.658 | 0.166 | -3.962 | 2,902.379 | 3.8e-05*** |
| **probRange33to066:noiseTypehighNoise** | **0.033** | **0.233** | **0.143** | **2,902.308** | **0.44327** |

#### Table 2.1.3: Experiment 2 (Second Added Boundary)

Dependent variable: certaintyEquivalent. Analysis Bin: 0.67 to 0.77.
Reference levels: probRange = '00to100', noiseType = 'lowNoise'.

| term | estimate | std.error | t-statistic | df | p.value |
|---|---|---|---|---|---|
| (Intercept) | -0.088 | 1.554 | -0.056 | 2,693.514 | 0.4775 |
| p | 48.683 | 2.109 | 23.085 | 2,655.069 | < 2e-16*** |
| probRange66to100 | 0.758 | 0.164 | 4.613 | 2,655.056 | 2.08e-06*** |
| noiseTypehighNoise | -0.741 | 0.187 | -3.969 | 2,655.283 | 3.71e-05*** |
| **probRange66to100:noiseTypehighNoise** | **0.575** | **0.262** | **2.193** | **2,655.172** | **0.0142*** |

#### Table 2.1.4: Experiment 3 (First Added Boundary)

Dependent variable: probabilityEquivalent. Analysis Bin: 0.35 to 0.45.
Reference levels: probRange = '00to100', noiseType = 'lowNoise'.

| term | estimate | std.error | t-statistic | df | p.value |
|---|---|---|---|---|---|
| (Intercept) | -4.953 | 1.340 | -3.697 | 3,867.026 | 0.000111*** |
| p | 112.787 | 3.259 | 34.608 | 3,839.033 | < 2e-16*** |
| probRange33to066 | -0.073 | 0.541 | -0.135 | 3,837.123 | 0.446162 |
| noiseTypehighNoise | -4.522 | 0.416 | -10.866 | 3,837.989 | < 2e-16*** |

| Dependent variable: probabilityEquivalent. Analysis Bin: 0.35 to 0.45.<br>Reference levels: probRange = '00to100', noiseType = 'lowNoise'. | | | | | |
|---|---|---|---|---|---|
| term | estimate | std.error | t-statistic | df | p.value |
| **probRange33to066:noiseTypehighNoise** | **4.193** | **0.587** | **7.143** | **3,837.776** | **5.45e-13*** ** |

Table 2.1.5: Experiment 3 (Second Added Boundary)

| Dependent variable: probabilityEquivalent. Analysis Bin: 0.67 to 0.77.<br>Reference levels: probRange = '00to100', noiseType = 'lowNoise'. | | | | | |
|---|---|---|---|---|---|
| term | estimate | std.error | t-statistic | df | p.value |
| (Intercept) | -32.404 | 3.727 | -8.694 | 2,873.568 | < 2e-16*** |
| p | 144.967 | 5.134 | 28.236 | 2,841.256 | < 2e-16*** |
| probRange66to100 | 0.028 | 0.545 | 0.052 | 2,840.116 | 0.479 |
| noiseTypehighNoise | -2.022 | 0.444 | -4.550 | 2,840.880 | 2.79e-06*** |
| **probRange66to100:noiseTypehighNoise** | **3.744** | **0.627** | **5.967** | **2,840.565** | **1.36e-09*** ** |

#### Underestimation Below Contextually Induced Boundaries with Added Noise

Tables 2.2.1 through 2.2.5 show that introducing new contextual boundaries in probability space leads to a systematic amplification of underestimation for probabilities just below those boundaries, under conditions of high cognitive noise when compared to when no boundaries are present there and low cognitive noise. As predicted, this effect is reflected in negative regression coefficients for the interaction term probRange:noiseTypeHighNoise. The observed effect is consistent with the hypothesis that cognitive noise interacts with newly induced structure in the probability space to enhance distortions in subjective probability representation.

##### Table 2.2.1: Experiment 1

Dependent variable: certaintyEquivalent. Analysis Bin: 0.40 to 0.50.
Reference levels: probRange = '00to100', noiseType = 'lowNoise'.

| term | estimate | std.error | t-statistic | df | p.value |
|---|---|---|---|---|---|
| (Intercept) | -1.904 | 1.243 | -1.532 | 1,792.010 | 0.0629. |
| p | 52.722 | 2.549 | 20.686 | 2,658.170 | < 2e-16*** |
| probRange00to050 | -0.289 | 0.203 | -1.425 | 2,658.135 | 0.0772. |
| noiseTypehighNoise | -1.394 | 0.204 | -6.835 | 2,658.181 | 5.07e-12*** |
| **probRange00to050:noiseTypehighNoise** | **-1.662** | **0.290** | **-5.739** | **2,658.299** | **5.31e-09*** ** |

##### Table 2.2.2: Experiment 2 (First Added Boundary)

Dependent variable: certaintyEquivalent. Analysis Bin: 0.23 to 0.33.
Reference levels: probRange = '00to100', noiseType = 'lowNoise'.

| term | estimate | std.error | t-statistic | df | p.value |
|---|---|---|---|---|---|
| (Intercept) | 2.302 | 0.556 | 4.142 | 1,479.979 | 1.82e-05*** |
| p | 45.117 | 1.774 | 25.438 | 2,922.257 | < 2e-16*** |
| probRange00to033 | -0.673 | 0.154 | -4.382 | 2,922.203 | 6.09e-06*** |
| noiseTypehighNoise | 0.559 | 0.164 | 3.418 | 2,922.381 | 0.00032*** |
| **probRange00to033:noiseTypehighNoise** | **-1.706** | **0.230** | **-7.424** | **2,922.429** | **7.42e-14*** ** |

##### Table 2.2.3: Experiment 2 (Second Added Boundary)

Dependent variable: certaintyEquivalent. Analysis Bin: 0.55 to 0.65.
Reference levels: probRange = '00to100', noiseType = 'lowNoise'.

| term | estimate | std.error | t-statistic | df | p.value |
|---|---|---|---|---|---|
| (Intercept) | -3.497 | 1.310 | -2.669 | 2,652.280 | 0.00383** |
| p | 54.485 | 2.125 | 25.642 | 2,640.250 | < 2e-16*** |
| probRange33to066 | 0.004 | 0.164 | 0.023 | 2,640.229 | 0.49072 |
| noiseTypehighNoise | -0.478 | 0.187 | -2.556 | 2,640.523 | 0.00532** |

Dependent variable: certaintyEquivalent. Analysis Bin: 0.55 to 0.65.
Reference levels: probRange = '00to100', noiseType = 'lowNoise'.

| term | estimate | std.error | t-statistic | df | p.value |
|---|---|---|---|---|---|
| **probRange33to066:noiseTypehighNoise** | **-0.704** | **0.263** | **-2.675** | **2,640.375** | **0.00376**** |

Table 2.2.4: Experiment 3 (First Added Boundary)

Dependent variable: probabilityEquivalent. Analysis Bin: 0.23 to 0.33.
Reference levels: probRange = '00to100', noiseType = 'lowNoise'.

| term | estimate | std.error | t-statistic | df | p.value |
|---|---|---|---|---|---|
| (Intercept) | 3.347 | 1.149 | 2.913 | 3,343.327 | 0.0018** |
| p | 88.045 | 3.839 | 22.936 | 3,368.029 | < 2e-16*** |
| probRange00to033 | -0.002 | 0.514 | -0.003 | 3,363.108 | 0.4988 |
| noiseTypehighNoise | 0.079 | 0.409 | 0.193 | 3,364.136 | 0.4235 |
| **probRange00to033:noiseTypehighNoise** | **-3.472** | **0.577** | **-6.020** | **3,363.942** | **9.64e-10***** |

Table 2.2.5: Experiment 3 (Second Added Boundary)

Dependent variable: probabilityEquivalent. Analysis Bin: 0.55 to 0.65.
Reference levels: probRange = '00to100', noiseType = 'lowNoise'.

| term | estimate | std.error | t-statistic | df | p.value |
|---|---|---|---|---|---|
| (Intercept) | -5.483 | 3.264 | -1.680 | 2,819.074 | 0.0465* |
| p | 109.091 | 5.359 | 20.358 | 2,763.215 | < 2e-16*** |
| probRange33to066 | 0.031 | 0.559 | 0.056 | 2,762.655 | 0.4778 |
| noiseTypehighNoise | -1.830 | 0.459 | -3.990 | 2,764.129 | 3.38e-05*** |
| **probRange33to066:noiseTypehighNoise** | **-3.145** | **0.646** | **-4.870** | **2,763.431** | **5.89e-07***** |

#### Inducing New Boundaries Creates New Probability Distortion Patterns (Main Effects of Boundaries for High Cognitive Noise)

The following results further test Prediction 2 by examining whether the presence of newly introduced contextual boundaries in probability space alters subjective probability distortions, even when cognitive noise is held constant at a high level. This analysis isolates the main effect of the probability range (probRange) under high-noise conditions, using trials that are matched in terms of objective probabilities and lotteries. In accordance with our preregistration, we fitted linear mixed-effects models with participant included as a random effect. Each model included fixed effects for the intercept, probability (p), noise type (low vs. high), probability range (reflecting whether a contextual boundary was introduced), and their interaction. To isolate the main effect of boundaries, we examined the probRange term specifically in the high-noise condition. As before, we focused on targeted bins of interest—namely, probability regions just above $p = 0.5$ in Experiment 1, and just above $p = 0.34$ and $p = 0.66$ in Experiments 2 and 3. We predicted that introducing boundaries will lead to changed distortions, resulting in overestimation of probabilities just above, and underestimation just below these new bounds. This would be reflected in a positive regression coefficient above introduced boundaries and negative regression coefficient below due to the contextual boundary manipulation alone. As in Prediction 1, Experiments 1 and 2 involved risky lottery valuation tasks and analyses were conducted on certainty equivalents (CE), while Experiment 3 involved judgments of explicit probability fractions and analyses were conducted on probability equivalents (PE).

##### Overestimation Above Contextually Induced Boundaries for High Noise

Tables 3.1.1 through 3.1.5 show that introducing new contextual boundaries in the probability space leads to a systematic overestimation for probabilities just above those boundaries, under conditions of high cognitive noise when compared to when no boundaries are present there. As predicted, this effect is reflected in positive regression coefficients for the term probRange. The observed distortions are consistent with the hypothesis that boundaries distorts subjective probability representation.

Table 3.1.1: Experiment 1

Dependent variable: certaintyEquivalent. Analysis Bin: 0.50 to 0.60.
Reference levels: probRange = '00to100', noiseType = 'highNoise'.

| term | estimate | std.error | t-statistic | df | p.value |
|---|---|---|---|---|---|
| (Intercept) | 9.936 | 1.349 | 7.366 | 1,941.891 | 1.29e-13*** |
| p | 28.933 | 2.286 | 12.654 | 2,685.128 | < 2e-16*** |
| **probRange50to100** | **1.701** | **0.181** | **9.377** | **2,685.152** | **< 2e-16*** ** |
| noiseTypelowNoise | -0.048 | 0.180 | -0.268 | 2,685.107 | 0.395 |
| probRange50to100:noiseTypelowNoise | -1.471 | 0.255 | -5.763 | 2,685.127 | 4.60e-09*** |

Table 3.1.2: Experiment 2 (First Added Boundary)

Dependent variable: certaintyEquivalent. Analysis Bin: 0.35 to 0.45.
Reference levels: probRange = '00to100', noiseType = 'highNoise'.

| term | estimate | std.error | t-statistic | df | p.value |
|---|---|---|---|---|---|
| (Intercept) | 1.238 | 0.805 | 1.538 | 2,163.489 | 0.0621. |
| p | 45.441 | 1.875 | 24.237 | 2,902.233 | < 2e-16*** |
| **probRange33to066** | **0.302** | **0.157** | **1.921** | **2,902.358** | **0.0274*** |
| noiseTypelowNoise | 0.658 | 0.166 | 3.962 | 2,902.379 | 3.8e-05*** |
| probRange33to066:noiseTypelowNoise | -0.033 | 0.233 | -0.143 | 2,902.308 | 0.4433 |

Table 3.1.3: Experiment 2 (Second Added Boundary)

Dependent variable: certaintyEquivalent. Analysis Bin: 0.67 to 0.77.
Reference levels: probRange = '00to100', noiseType = 'highNoise'.

| term | estimate | std.error | t-statistic | df | p.value |
|---|---|---|---|---|---|
| (Intercept) | -0.829 | 1.557 | -0.532 | 2,694.116 | 0.2973 |
| p | 48.683 | 2.109 | 23.085 | 2,655.069 | < 2e-16*** |
| **probRange66to100** | **1.332** | **0.204** | **6.528** | **2,655.258** | **3.97e-11*** ** |
| noiseTypelowNoise | 0.741 | 0.187 | 3.969 | 2,655.283 | 3.71e-05*** |

| Dependent variable: certaintyEquivalent. Analysis Bin: 0.67 to 0.77.<br>Reference levels: probRange = '00to100', noiseType = 'highNoise'. | | | | | |
|---|---|---|---|---|---|
| term | estimate | std.error | t-statistic | df | p.value |
| probRange66to100:noiseType lowNoise | -0.575 | 0.262 | -2.193 | 2,655.172 | 0.0142* |

Table 3.1.4: Experiment 3 (First Added Boundary)

| Dependent variable: probabilityEquivalent. Analysis Bin: 0.35 to 0.45.<br>Reference levels: probRange = '00to100', noiseType = 'highNoise'. | | | | | |
|---|---|---|---|---|---|
| term | estimate | std.error | t-statistic | df | p.value |
| (Intercept) | -9.475 | 1.329 | -7.131 | 3,864.334 | 5.94e-13*** |
| p | 112.787 | 3.259 | 34.608 | 3,839.033 | < 2e-16*** |
| **probRange33to066** | **4.119** | **0.228** | **18.062** | **3,841.247** | **< 2e-16*** ** |
| noiseTypelowNoise | 4.522 | 0.416 | 10.866 | 3,837.989 | < 2e-16*** |
| probRange33to066:noiseType lowNoise | -4.193 | 0.587 | -7.143 | 3,837.776 | 5.45e-13*** |

Table 3.1.5: Experiment 3 (Second Added Boundary)

| Dependent variable: probabilityEquivalent. Analysis Bin: 0.67 to 0.77.<br>Reference levels: probRange = '00to100', noiseType = 'highNoise'. | | | | | |
|---|---|---|---|---|---|
| term | estimate | std.error | t-statistic | df | p.value |
| (Intercept) | -34.427 | 3.716 | -9.265 | 2,873.783 | < 2e-16*** |
| p | 144.967 | 5.134 | 28.236 | 2,841.255 | < 2e-16*** |
| **probRange66to100** | **3.772** | **0.310** | **12.166** | **2,841.947** | **< 2e-16*** ** |
| noiseTypelowNoise | 2.022 | 0.444 | 4.550 | 2,840.880 | 2.79e-06*** |
| probRange66to100:noiseType lowNoise | -3.744 | 0.627 | -5.967 | 2,840.565 | 1.36e-09*** |

#### Underestimation Below Contextually Induced Boundaries for High Noise

Tables 3.2.1 through 3.2.5 show that introducing new contextual boundaries in the probability space leads to a systematic underestimation for probabilities just below those boundaries, under conditions of high cognitive noise when compared to when no boundaries are present there. As predicted, this effect is reflected in negative regression coefficients for the term probRange. The observed distortions are consistent with the hypothesis that boundaries distortions subjective probability representation.

#### Table 3.2.1: Experiment 1

Dependent variable: certaintyEquivalent. Analysis Bin: 0.40 to 0.50.
Reference levels: probRange = '00to100', noiseType = 'highNoise'.

| term | estimate | std.error | t-statistic | df | p.value |
|---|---|---|---|---|---|
| (Intercept) | -3.298 | 1.244 | -2.652 | 1,792.717 | 0.00403** |
| p | 52.722 | 2.549 | 20.686 | 2,658.170 | < 2e-16*** |
| **probRange00to050** | **-1.952** | **0.207** | **-9.447** | **2,658.406** | **< 2e-16*** ** |
| noiseTypelowNoise | 1.394 | 0.204 | 6.835 | 2,658.181 | 5.07e-12*** |
| probRange00to050:noiseTypelowNoise | 1.662 | 0.290 | 5.739 | 2,658.299 | 5.31e-09*** |

#### Table 3.2.2: Experiment 2 (First Added Boundary)

Dependent variable: certaintyEquivalent. Analysis Bin: 0.23 to 0.33.
Reference levels: probRange = '00to100', noiseType = 'highNoise'.

| term | estimate | std.error | t-statistic | df | p.value |
|---|---|---|---|---|---|
| (Intercept) | 2.860 | 0.545 | 5.251 | 1,411.461 | 8.70e-08*** |
| p | 45.117 | 1.774 | 25.438 | 2,922.257 | < 2e-16*** |
| **probRange00to033** | **-2.379** | **0.171** | **-13.923** | **2,922.579** | **< 2e-16*** ** |
| noiseTypelowNoise | -0.559 | 0.164 | -3.418 | 2,922.381 | 0.00032*** |
| probRange00to033:noiseTypelowNoise | 1.706 | 0.230 | 7.424 | 2,922.429 | 7.42e-14*** |

217 Table 3.2.3: Experiment 2 (Second Added Boundary)

Dependent variable: certaintyEquivalent. Analysis Bin: 0.55 to 0.65.
Reference levels: probRange = '00to100', noiseType = 'highNoise'.

| term | estimate | std.error | t-statistic | df | p.value |
|---|---|---|---|---|---|
| (Intercept) | -3.974 | 1.309 | -3.036 | 2,651.819 | 0.001209** |
| p | 54.485 | 2.125 | 25.642 | 2,640.250 | < 2e-16*** |
| **probRange33to066** | **-0.700** | **0.206** | **-3.406** | **2,640.470** | **0.000335*** ** |
| noiseTypelowNoise | 0.478 | 0.187 | 2.556 | 2,640.523 | 0.005319** |
| probRange33to066:noiseType lowNoise | 0.704 | 0.263 | 2.675 | 2,640.375 | 0.003765** |

218 Table 3.2.4: Experiment 3 (First Added Boundary)

Dependent variable: probabilityEquivalent. Analysis Bin: 0.23 to 0.33.
Reference levels: probRange = '00to100', noiseType = 'highNoise'.

| term | estimate | std.error | t-statistic | df | p.value |
|---|---|---|---|---|---|
| (Intercept) | 3.426 | 1.072 | 3.195 | 3,296.826 | 0.000706*** |
| p | 88.045 | 3.839 | 22.936 | 3,368.029 | < 2e-16*** |
| **probRange00to033** | **-3.474** | **0.262** | **-13.251** | **3,366.981** | **< 2e-16*** ** |
| noiseTypelowNoise | -0.079 | 0.409 | -0.193 | 3,364.136 | 0.423458 |
| probRange00to033:noiseType lowNoise | 3.472 | 0.577 | 6.020 | 3,363.942 | 9.64e-10*** |

219 Table 3.2.5: Experiment 3 (Second Added Boundary)

Dependent variable: probabilityEquivalent. Analysis Bin: 0.55 to 0.65.
Reference levels: probRange = '00to100', noiseType = 'highNoise'.

| term | estimate | std.error | t-statistic | df | p.value |
|---|---|---|---|---|---|
| (Intercept) | -7.313 | 3.238 | -2.258 | 2,819.346 | 0.012* |
| p | 109.091 | 5.359 | 20.358 | 2,763.215 | < 2e-16*** |
| **probRange33to066** | **-3.114** | **0.324** | **-9.615** | **2,765.710** | **< 2e-16*** ** |
| noiseTypelowNoise | 1.830 | 0.459 | 3.990 | 2,764.129 | 3.38e-05*** |

Dependent variable: probabilityEquivalent. Analysis Bin: 0.55 to 0.65.
Reference levels: probRange = '00to100', noiseType = 'highNoise'.

| term | estimate | std.error | t-statistic | df | p.value |
|---|---|---|---|---|---|
| probRange33to066:noiseType lowNoise | 3.145 | 0.646 | 4.870 | 2,763.431 | 5.89e-07*** |

#### Inducing New Boundaries Reduces Variability in Lottery Valuation and Probability Estimation Close to the Boundaries

The following tables show the results of analyses testing Prediction 3, which proposes that introducing new contextual boundaries reduces behavioral variability close to the newly introduced boundaries, particularly under conditions of high cognitive noise. Unlike Predictions 1 and 2, which focus on mean distortions in subjective probability weighting, Prediction 3 addresses the prediction that contextual boundary structure can suppress variability in responses. To quantify behavioral variability, as per our pre-pregistered analysis pipeline, we first computed each participant's trial-wise deviations from their mean response (certainty equivalents in Experiments 1 and 2, probability equivalents in Experiment 3) at each probability level given by (d(p) = $\frac{1}{n}\sum_i |s_i - \frac{1}{n}\sum_j s_j|$). These absolute deviations were then averaged across trials to obtain a single variability measure per probability level. Finally, as per our pre-registrations, we applied a rolling mean over 5 adjacent probability levels (corresponding to roughly 0.10 in probability space): (var(p) = $\frac{1}{5}\sum_{\{k=-2\}}^{\{k=2\}} d(p+k\Delta p)$, where $\Delta p$ is 0.02 probability. This rolling average of variability provided a smoothed out localized measure of behavioral variability (variability_rolling_avg), capturing how much responses fluctuated. We then tested whether the introduction of contextual boundaries (e.g., at p = 0.5 in Experiment 1, and p = 0.34 and p = 0.66 in Experiments 2 and 3) led to reduced variability just above and below these boundaries (in bins of size 0.15), in the high noise condition. Linear mixed-effects models were fitted within these bins, with variability_rolling_avg as the dependent variable, and fixed effects for probability (p), probRange, noiseType and the interaction of probRange and noiseType. Subject was included as a random effect. The key prediction was a **negative effect of probRange** (boundary vs. no-boundary) on behavioral variability both below and above introduced boundaries.

#### Reduced Behavioral Variability Above Contextually Induced Boundaries for High Noise

Tables 4.1.1 through 4.1.5 show that introducing new contextual boundaries in the probability space leads to a systematic decrease in behavioral variability just above those boundaries, due to suppression in cognitive noise due to those boundaries. As predicted, this effect is reflected in negative regression coefficients for the term probRange.

##### Table 4.1.1: Experiment 1

Dependent variable: rolling_variability. Analysis Bin: 0.5 to 0.65.
Reference levels: probRange = '00to100', noiseType = 'highNoise'.

| term | estimate | std.error | t-statistic | df | p.value |
|---|---|---|---|---|---|
| (Intercept) | -0.409 | 0.219 | -1.863 | 2,991.931 | 0.0312* |
| p | 3.358 | 0.363 | 9.242 | 4,062.207 | < 2e-16*** |
| **probRange50to100** | **-0.308** | **0.046** | **-6.680** | **4,062.476** | **1.35e-11*** ** |
| noiseTypelowNoise | -0.633 | 0.044 | -14.221 | 4,062.238 | < 2e-16*** |
| probRange50to100:noiseType lowNoise | 0.081 | 0.063 | 1.290 | 4,062.295 | 0.0986. |

##### Table 4.1.2: Experiment 2 (First Added Boundary)

Dependent variable: rolling_variability. Analysis Bin: 0.35 to 0.5.
Reference levels: probRange = '00to100', noiseType = 'highNoise'.

| term | estimate | std.error | t-statistic | df | p.value |
|---|---|---|---|---|---|
| (Intercept) | 1.733 | 0.150 | 11.550 | 999.2524 | < 2e-16*** |
| p | -0.221 | 0.305 | -0.725 | 4,268.1191 | 0.234 |
| **probRange33to066** | **-0.440** | **0.039** | **-11.161** | **4,268.2470** | **< 2e-16*** ** |
| noiseTypelowNoise | -0.823 | 0.039 | -20.908 | 4,268.1828 | < 2e-16*** |
| probRange33to066:noiseType lowNoise | 0.218 | 0.056 | 3.926 | 4,268.1763 | 4.38e-05*** |

Table 4.1.3: Experiment 2 (Second Added Boundary)

Dependent variable: rolling_variability. Analysis Bin: 0.67 to 0.82.
Reference levels: probRange = '00to100', noiseType = 'highNoise'.

| term | estimate | std.error | t-statistic | df | p.value |
|---|---|---|---|---|---|
| (Intercept) | 1.285 | 0.264 | 4.866 | 3,523.142 | 5.94e-07*** |
| p | 0.492 | 0.337 | 1.459 | 4,030.085 | 0.0724. |
| **probRange66to100** | **-0.361** | **0.043** | **-8.357** | **4,030.142** | **< 2e-16*** ** |
| noiseTypelowNoise | -0.729 | 0.042 | -17.349 | 4,030.169 | < 2e-16*** |
| probRange66to100:noiseType lowNoise | 0.086 | 0.059 | 1.463 | 4,030.078 | 0.0718. |

Table 4.1.4: Experiment 3 (First Added Boundary)

Dependent variable: rolling_variability. Analysis Bin: 0.35 to 0.5.
Reference levels: probRange = '00to100', noiseType = 'highNoise'.

| term | estimate | std.error | t-statistic | df | p.value |
|---|---|---|---|---|---|
| (Intercept) | 1.121 | 0.292 | 3.841 | 765.5285 | 6.64e-05*** |
| p | 7.165 | 0.592 | 12.094 | 5,247.3859 | < 2e-16*** |
| **probRange33to066** | **-1.338** | **0.059** | **-22.747** | **5,247.7762** | **< 2e-16*** ** |
| noiseTypelowNoise | -4.131 | 0.099 | -41.835 | 5,247.2545 | < 2e-16*** |
| probRange33to066:noiseType lowNoise | 1.341 | 0.140 | 9.610 | 5,247.1820 | < 2e-16*** |

Table 4.1.5: Experiment 3 (Second Added Boundary)

Dependent variable: rolling_variability. Analysis Bin: 0.67 to 0.82.
Reference levels: probRange = '00to100', noiseType = 'highNoise'.

| term | estimate | std.error | t-statistic | df | p.value |
|---|---|---|---|---|---|
| (Intercept) | 1.197 | 0.548 | 2.184 | 3,530.842 | 0.0145* |
| p | 4.206 | 0.700 | 6.010 | 4,735.289 | 9.96e-10*** |
| **probRange66to100** | **-1.222** | **0.065** | **-18.823** | **4,735.424** | **< 2e-16*** ** |
| noiseTypelowNoise | -4.310 | 0.104 | -41.513 | 4,735.242 | < 2e-16*** |

Dependent variable: rolling_variability. Analysis Bin: 0.67 to 0.82.
Reference levels: probRange = '00to100', noiseType = 'highNoise'.

| term | estimate | std.error | t-statistic | df | p.value |
|---|---|---|---|---|---|
| probRange66to100:noiseType lowNoise | 1.222 | 0.146 | 8.354 | 4,735.064 | < 2e-16*** |

#### Reduced Behavioral Variability Below Contextually Induced Boundaries for High Noise

Tables 4.2.1 through 4.2.5 show that introducing new contextual boundaries in the probability space leads to a systematic decrease in behavioral variability just below those boundaries as well, due to suppression in cognitive noise due to those boundaries. As predicted, this effect is reflected in negative regression coefficients for the term probRange.

##### Table 4.2.1: Experiment 1

Dependent variable: rolling_variability. Analysis Bin: 0.35 to 0.50.
Reference levels: probRange = '00to100', noiseType = 'highNoise'.

| term | estimate | std.error | t-statistic | df | p.value |
|---|---|---|---|---|---|
| (Intercept) | 2.332 | 0.184 | 12.652 | 773.009 | < 2e-16*** |
| p | -1.196 | 0.364 | -3.287 | 4,292.151 | 0.000511*** |
| **probRange00to050** | **-0.294** | **0.047** | **-6.222** | **4,292.444** | **2.7e-10*** ** |
| noiseTypelowNoise | -0.883 | 0.047 | -18.906 | 4,292.211 | < 2e-16*** |
| probRange00to050:noiseType lowNoise | 0.235 | 0.066 | 3.548 | 4,292.296 | 0.000196*** |

##### Table 4.2.2: Experiment 2 (First Added Boundary)

Dependent variable: rolling_variability. Analysis Bin: 0.18 to 0.33.
Reference levels: probRange = '00to100', noiseType = 'highNoise'.

| term | estimate | std.error | t-statistic | df | p.value |
|---|---|---|---|---|---|
| (Intercept) | 1.442 | 0.102 | 14.101 | 410.0912 | < 2e-16*** |

| Dependent variable: rolling_variability. Analysis Bin: 0.18 to 0.33.<br>Reference levels: probRange = '00to100', noiseType = 'highNoise'. | | | | | |
|---|---|---|---|---|---|
| term | estimate | std.error | t-statistic | df | p.value |
| p | -0.543 | 0.300 | -1.810 | 4,015.1999 | 0.03515* |
| **probRange00to033** | **-0.101** | **0.039** | **-2.608** | **4,015.3929** | **0.00457**** |
| noiseTypelowNoise | -0.485 | 0.037 | -13.002 | 4,015.1868 | < 2e-16*** |
| probRange00to033:noiseType lowNoise | -0.081 | 0.053 | -1.543 | 4,015.2377 | 0.06144. |

Table 4.2.3: Experiment 2 (Second Added Boundary)

| Dependent variable: rolling_variability. Analysis Bin: 0.51 to 0.66.<br>Reference levels: probRange = '00to100', noiseType = 'highNoise'. | | | | | |
|---|---|---|---|---|---|
| term | estimate | std.error | t-statistic | df | p.value |
| (Intercept) | -0.269 | 0.194 | -1.391 | 1,927.130 | 0.0822. |
| p | 2.919 | 0.311 | 9.397 | 4,011.124 | < 2e-16*** |
| **probRange33to066** | **-0.199** | **0.039** | **-5.059** | **4,011.152** | **2.2e-07***** |
| noiseTypelowNoise | -0.646 | 0.038 | -16.930 | 4,011.193 | < 2e-16*** |
| probRange33to066:noiseType lowNoise | 0.020 | 0.054 | 0.366 | 4,011.115 | 0.3572 |

Table 4.2.4: Experiment 3 (First Added Boundary)

| Dependent variable: rolling_variability. Analysis Bin: 0.18 to 0.33.<br>Reference levels: probRange = '00to100', noiseType = 'highNoise'. | | | | | |
|---|---|---|---|---|---|
| term | estimate | std.error | t-statistic | df | p.value |
| (Intercept) | 5.943 | 0.253 | 23.500 | 391.104 | < 2e-16*** |
| p | -7.585 | 0.776 | -9.771 | 4,654.709 | < 2e-16*** |
| **probRange00to033** | **-0.691** | **0.072** | **-9.544** | **4,654.693** | **< 2e-16***** |
| noiseTypelowNoise | -3.971 | 0.115 | -34.637 | 4,654.155 | < 2e-16*** |
| probRange00to033:noiseType lowNoise | 0.690 | 0.162 | 4.263 | 4,654.134 | 1.03e-05*** |

##### Table 4.2.5: Experiment 3 (Second Added Boundary)

Dependent variable: rolling_variability. Analysis Bin: 0.51 to 0.66.
Reference levels: probRange = '00to100', noiseType = 'highNoise'.

| term | estimate | std.error | t-statistic | df | p.value |
|---|---|---|---|---|---|
| (Intercept) | -1.583 | 0.471 | -3.358 | 1,862.704 | 4e-04*** |
| p | 10.533 | 0.763 | 13.802 | 4,654.220 | < 2e-16*** |
| **probRange33to066** | **-1.001** | **0.070** | **-14.361** | **4,654.563** | **< 2e-16*** ** |
| noiseTypelowNoise | -4.526 | 0.111 | -40.880 | 4,654.236 | < 2e-16*** |
| probRange33to066:noiseTypelowNoise | 1.001 | 0.156 | 6.431 | 4,654.080 | 6.98e-11*** |

### Supplementary Note 5: Source of Contextual Boundary Repulsions

To determine whether contextual boundary repulsions originate during adaptive encoding or Bayesian decoding, we examined whether noise-driven biases and variability are altered in regions far away from the contextually induced boundaries. This test leverages a key theoretical distinction: Under efficient encoding, representational resources adapt to the input range. As a result, the same internal noise corresponds to different ranges of inputs and thereby, any existing noise-dependent distortions and variance amplify as a function of the range. If encoding adapts to a new, narrower contextual range, we should observe corresponding reductions in both bias and variability even at distant points (near 0 and 1). In contrast, under non-adaptive encoding, boundary repulsions arise simply from decoding. In this case the prior impacts the posterior during decoding, but encoding remains unchanged and the effects of the new contextual boundary remain local, with no change in biases and variability far away from the induced boundary. As a result, biases and variability at the natural boundaries of 0 and 1 do not change as a function of the range, indicating that the newly generated biases and variance at the new boundaries are decoding-based in origin.

To test this, we conducted exploratory Bayes Factor model comparisons evaluating whether including the probRange variable improved model fits for bias (certainty/probability equivalents) and variability in regions far from the contextual boundary (i.e., near 0 and 1). Specifically, we analyzed behavior within two bins: (0–0.1) and (0.9–1.0). These bins were chosen because they lie far from the contextually induced boundary, minimizing the boundary repulsion effect due to the induced boundaries themselves. This separation was crucial to disentangle changes due to contextual boundaries from range-dependent effects far from the contextually induced boundaries. For bins in the experimental condition (Risk Single, Risk Double, and No Risk), we computed Bayes Factors comparing the full model with predictors p, probRange, and subject (random) and the reduced model excluding probRange (i.e., only p and subject). The resulting Bayes Factors ($BF_{10}$) quantify the strength of evidence in favor of including probRange as a predictor. Values above 1 favor the full model; values below 1 favor the reduced model. Therefore, values above 1 suggest that there is a range-dependent change in the variable of interest (bias or variability) even far away from the induced boundaries, in the bins near the natural probability boundaries.

The results (see Table 5) showed that, in nearly all conditions and bins, $BF_{10}$ values were well below 1, indicating no evidence for range-dependent changes in bias or variability near the natural boundaries. This pattern supports the view that contextual boundary repulsions primarily reflect Bayesian decoding mechanisms, where prior truncation near the boundary drives local distortions without altering distant regions of the stimulus space. A notable exception was found in Experiment 2 (Risk Double) within the [0–0.1] bin. In this case, both bias and variability showed strong evidence for range-dependent changes. This convergence of effects in both measures suggests a potential case of adaptive encoding in this particular condition, but a decoding-based origin everywhere else. Taken together, these results indicate that the repulsions observed in our task do not systematically extend to distant regions of the input space. Instead, they are localized to the contextual boundaries and are best explained by Bayesian decoding of fixed, non-adaptive internal representations.

#### Table 5: Effects far away from the contextually induced boundaries

| Experiment | BF10 | Error | Bin | Type |
|---|---|---|---|---|
| Exp 1: Risk Single | 1.166836e-01 | 0.0432835 | 0–0.10 | Bias/Distortion (Above 0) |

| Experiment | BF10 | Error | Bin | Type |
|---|---|---|---|---|
| Exp 2: Risk Double | 3.928399e+17 | 0.0404260 | 0–0.10 | Bias/Distortion (Above 0) |
| Exp 3: No Risk | 1.843110e-01 | 0.3097847 | 0–0.10 | Bias/Distortion (Above 0) |
| Exp 1: Risk Single | 2.333715e-01 | 0.1433492 | 0.90–1 | Bias/Distortion (Below 1) |
| Exp 2: Risk Double | 6.232790e-02 | 0.0459257 | 0.90–1 | Bias/Distortion (Below 1) |
| Exp 3: No Risk | 3.304990e-02 | 0.1246364 | 0.90–1 | Bias/Distortion (Below 1) |
| Exp 1: Risk Single | 5.702140e-02 | 0.1108605 | 0–0.10 | Variance (Above 0) |
| Exp 2: Risk Double | 4.522868e+02 | 0.1536471 | 0–0.10 | Variance (Above 0) |
| Exp 3: No Risk | 1.780499e-01 | 0.2308722 | 0–0.10 | Variance (Above 0) |
| Exp 1: Risk Single | 2.782169e-01 | 0.2434397 | 0.90–1 | Variance (Below 1) |
| Exp 2: Risk Double | 6.752890e-02 | 0.0705112 | 0.90–1 | Variance (Below 1) |
| Exp 3: No Risk | 1.769490e-02 | 0.2540840 | 0.90–1 | Variance (Below 1) |

### Supplementary Note 6: Instructions Used

#### Instructions For Risky Valuation Task

Below, we provide instructions used for the risky decision making experiment with one induced boundary at 0.5. The instructions for the two boundary condition were exactly the same except for the indication of the ranges.

### Instructions

Thank you for joining our study! Please read the following instructions carefully. You will then complete a short comprehension quiz at the end before you can proceed further in the study.

**Your task**

You will decide whether you want to participate in different lotteries that will be presented on the screen. Each lottery gives you a chance of winning 50 **Experimental Currency Units (ECU)** with different probabilities (p). For every round, the probability (p) on the screen represents your chance of winning 50 ECU, while the remaining probability (1-p) means you might not win anything (0 ECU). Simply put, the higher the probability shown, the more attractive the lottery, and the better your chances of winning the 50 ECU. Note that at the end of the experiment, the amount you win in ECU will be converted to CHF, with a conversion rate of 50 ECU to 40 CHF.

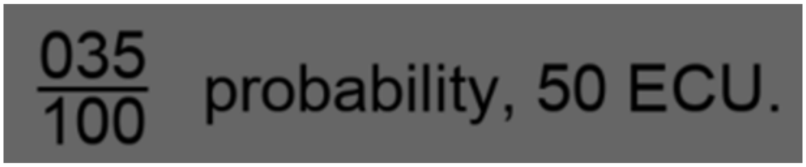


**Figure 1** Throughout the experiment, each round will present lotteries where you can potentially win a constant amount of 50 ECU. The probability of winning 50 ECU changes in each round.

Your task in each round today is to specify the minimum amount of money you would have to be paid for sure to not participate in the lottery. There are no right or wrong answers; your responses depend only on how valuable it is to you to participate in the lottery.

For example, if you indicate V ECU in response to the lottery in Figure 1, this is like saying:

**To me, receiving V ECU FOR SURE is as valuable as a 35% chance of winning 50 ECU. Any offer less than V ECU would not tempt me away from the lottery, but any offer V ECU or higher would make me choose this payoff and not the lottery.**

You will be paid based on six of the choices that you make throughout the experiment, which will be selected randomly in the end. The payment is implemented in a way that it is best for you to type the exact amount V that is as valuable to you as playing the lottery: Typing this amount ensures the best outcome for you in terms of whether you get to play the lottery or you get a payoff that is worth more to you than playing the lottery. Indicating any other amount may result in suboptimal outcomes for you. The exact mechanism - and the reason why you should carefully consider and report how much the lottery is worth to you - will be explained in the payment mechanism section at the end of the instructions.

**Important details of the experiment:**

**Different probability ranges**

The experiment has **two parts**, with different **ranges of probabilities**, as explained below. The order in which you do these two parts will be randomly selected. Before each part, you'll be informed of the specific range of probabilities that will be presented. For all parts, the probabilities will be determined randomly for every round, but will always lie within the specified range. The winning amount is always 50 ECU.

1) **Full range part**: The lotteries will pay 50 ECU, with probabilities ranging from 0.01 to 0.99 (the full range). These lotteries thus vary in their financial attractiveness from least attractive lottery (1% probability of winning 50 ECU) to the most attractive lottery (99% probability of winning 50 ECU).

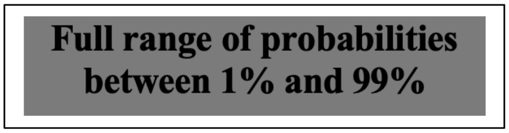


2) **Half range part**: The half range part of the experiment itself has two sub-sections that correspond to two different half ranges.
    a. In the **lower-half-range** sub-section, the lotteries will pay 50 ECU with probabilities in the restricted range between 0.01-0.49. In these blocks, you will never see a lottery offering a probability of winning higher than 49%. Therefore, lotteries in this sub-block are less financially attractive.

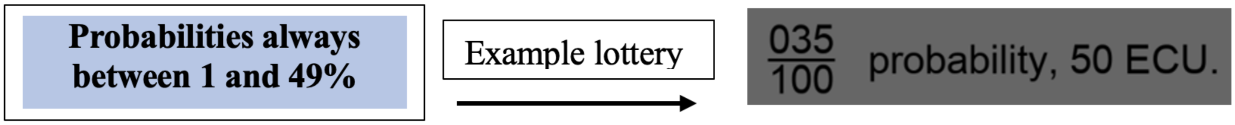


    b. In the **upper-half-range** part, the lotteries will pay 50 ECU with probabilities in the restricted range between 0.51-0.99. In these blocks, you will never see a lottery offering a probability of winning lower than 51%. Therefore, lotteries in this sub-block are more financially attractive.

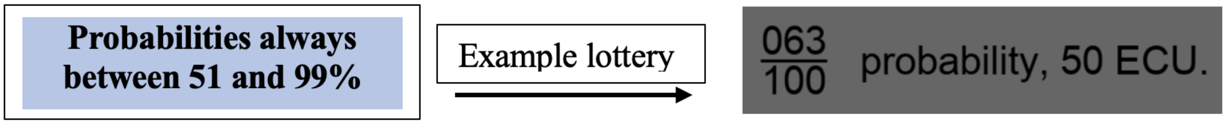


**Different complexity of payoff presentation**

In both parts of the experiment (full-range and half-range), there will be **two types of blocks** that differ in how **complex** it is to determine the probabilities.

1) In **Easy blocks,** probabilities are presented as easy fractions (x/y) with a fixed y of 100 -- only the x changes (see Figure 2A). The x/100 therefore directly represents the probability with which you can win the 50 ECU with.
2) In **hard blocks**, both the x and y in the fraction (x/y) are randomly selected 4-digit numbers, and both change in each round (see Figure 2B). You will thus have to estimate, by approximate division of x and y, the probability with which you can win the 50 ECU with.

a) 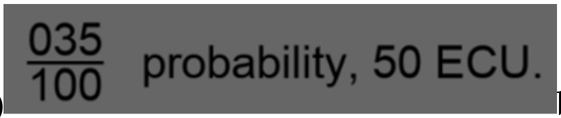

b) 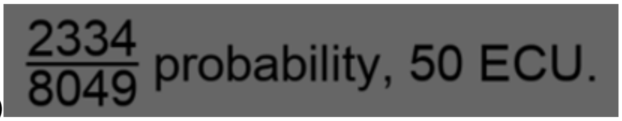


**Figure 2** A) Easy Blocks: Half of today's blocks present winning chances as fractions with a fixed denominator of 100. Only the numerator changes between rounds. B) Hard Blocks: The remaining half of the blocks will present the percentage chance of winning as complex 4-digit fractions, with both the numerator and the denominator changing between rounds.

It is difficult to approximate the payoff in the hard blocks, but this is part of the experiment. You are not permitted to use any type of calculator (if you are seen using a calculator, we will have to disqualify you from the experiment without any payment). Although it is easier to determine the probability with a denominator of 100 (Figure 2A), please try to make decisions as well as you can in both the easy and hard conditions.

**General procedure of the experiment**

Your task today is to indicate the minimum sure amount to opt out of playing the lottery in each round.

You have two main parts of the experiment, each with specific characteristics. In summary, you'll encounter:

1) Full range part: (**all probabilities between 1 and 99%)**. This part has 2 blocks –
    1) **Easy** probability presentation.
    2) **Hard** probability presentation.

2a) In the **lower-half-range part**, probabilities vary between **1 and 49%**. This part has two blocks:

3) **Easy** probability presentation
4) **Hard** probability presentation.

2b) In the **upper-half-range part**, probabilities vary between **51 and 99%**. This part has two blocks:

5) **Easy** probability presentation
6) **Hard** probability presentation.

#### Please note

- The order in which you will see the half and the full range part of the experiment is randomly determined. Within the half range part, whether you see the more attractive (upper half range) or less attractive lotteries (lower half range) is also determined randomly. So always pay attention at the start of a block to see in which condition you are in!
- The order of "easy" and "hard blocks" is also determined randomly.
- Pay close attention to the information provided before each block, which specifies the condition you'll encounter.

**Details on your response**

For every lottery, please input the sure amount V by typing numbers at the top of the keyboard and the decimal key, then please press <enter> to finalize your choice. Note that this is necessary to record your response. As you type, the numbers will be displayed in black; pressing <enter> will turn them green and indicate your final decision. After 0.5 seconds of your response, the next round will start, and you will be presented with the next lottery. Once you have submitted your response, you cannot change your input anymore so try to ensure that the valuation you entered is correct.

You have at most 9 seconds to indicate the sure amount V, ranging from 0 to 50 ECU. By using decimal values, you can specify any amount within this range. If you submit a number outside this range, then the answer won't be accepted and the input turns red. You can still modify your input if this happens (if your maximum time of 9 seconds is not over), by pressing <backspace>. You will be paid based on six choices you took, which are randomly drawn from all rounds you played. Importantly, the payment mechanism is designed in such a way that it will be optimal for you to carefully consider what price would be right for you to give up the lottery and indicate this price as precisely as possible.

**Payment Details**

As explained above, your task in each round is to specify the minimum amount of money V you would need to be paid for sure to opt out of the gamble. Later, the computer randomly selects 6 rounds and pays you according to your choice. Here's how it works:

1. **Computer makes a sure Offer S that is randomly determined:**
   - The computer offers a sure amount between 0 and 50 ECU for a selected round, which is randomly determined.

**2. Comparison of offered random amount S with the valuation V you indicated:**
   - If the offered amount S is below your valuation V:
     - You prefer the lottery and decline the sure offer (since the offer is lower than how much you value the lottery). The lottery is played and you get its outcome.
   - If the offered amount S is the same or above your valuation V:
     - You accept and receive the sure amount S offered by the computer. The lottery is not played.

This process ensures that your choices determine whether you play the lottery or accept the sure amount offered on each selected round.

**Frequently asked question: Can I get a better outcome by indicating a sure valuation V that is larger or smaller than my real preference?**

**Answer: No.** Indicating an amount different from your true preferences can result in you getting a suboptimal option. Let's explore scenarios:

**Scenario 1: Misreporting by indicating a smaller value V for the lottery**

- If you state 3 ECU is equivalent for you to a 40% chance of winning 50 ECU (while your true preference is 20 ECU), and the computer offers 3.4 ECU for sure:
  - The computer pays 3.4 ECU, and you miss the chance to play this lottery as it is inferred that you prefer it to the lottery.
  - You're worse off, receiving a lower sure amount S instead of the lottery you prefer.

**Scenario 2: Misreporting by indicating a Larger Value V for the lottery**

- If you state 40.3 ECU is equivalent to a 40% chance of winning 50 ECU (while your true preference is 20 ECU), and the computer offers 39.2 ECU for sure:
  - The computer makes you play the lottery instead of paying the offered 39.2 ECU, as it is inferred that you prefer the lottery.
  - You're worse off, playing the lottery instead of receiving the higher sure amount that you would actually prefer.

**Conclusion:** Misreporting means you can get suboptimal outcomes. So please report your true valuation for the displayed lottery carefully, to make sure that you actually get the option you prefer when the computer later makes a random offer. These decisions in the experiment will influence your overall payout.

At the end of the experiment, the computer randomly selects 6 rounds, one from each of the 6 conditions you play today. The 6 conditions you play today are the easy and hard conditions for the full range part (2), the easy and hard conditions for the 1-49% half range part (2), and the easy and hard conditions for the 51-99% half range part (2). The payoff will be based on the values you indicate for those lotteries, and the comparison with the sure offers the computer makes for these rounds. The average amount that you win in those 6 randomly selected rounds, based on your choices and the lottery outcomes, determines your bonus.

Winning the lottery in all 6 of those selected rounds would add an extra 50 ECU which is equivalent to an extra 40 CHF. Your total earnings, including the participation fee of 10 CHF, can thus range from 10 to 50 CHF. Keep in mind that if a round is randomly chosen for which you didn't indicate a value, you receive no bonus for that round. Make sure you respond timely and precisely (by clicking <enter>) for all rounds to maximize your payment. The total payout, including participation fees, will be displayed on the screen after the experiment.

**Summary**

1. **Valuation task:** In each round, specify the minimum amount V in ECU that you require to opt out of playing the lottery gamble. You have a **Time limit** of 9 seconds to type your answer and **submit it by pressing <enter>. You do not have to wait 9 seconds for the next lottery if you respond earlier.**
2. **Two parts** of this experiment: You are randomly assigned to complete one of them first and the other next.
    a. In one part of the experiment, you will complete 2 blocks with probabilities covering the full range from 1%-99%.
        i. One block will have "easy" probabilities.
        ii. The other block will have "hard" probabilities.
        iii. You will be randomly assigned to do one of these first.
    b. In another part of the experiment, you will complete 4 blocks where probabilities will be in a restricted range.
        i. You are randomly assigned to start with the 2 "hard" or the 2 "easy" blocks first.
        ii. For both easy and hard blocks, you will complete both the 1-49% restricted half range (less attractive lotteries) and the 51-99% restricted half range (more attractive lotteries) conditions, assigned in random order.

3. We implemented a payment mechanism that makes it optimal for you to indicate the sure amount that is worth as much to you as playing the lottery – not more and not less.

**4. Important reminder:** Always try to indicate on each round how much you value that lottery. If you fail to do so, or if you keep generating invalid responses (above 50 ECU) and that round is selected, you will not be given any bonus. This will bring down your total payment.

**5. Potential earnings:** These can range between 10 and 50 CHF, based on the rounds selected by the computer, your choices in those rounds, and the outcomes of the lotteries.
**6. Warning:** Do not use any calculator, or you will not be able to take part in the experiment and receive your payment.

If you have any further questions, you can call the experimenter and clarify these. If you do not have any further questions, you can begin answering the comprehension questions on the screen. From now on, you will only need the keyboard to complete the study.

#### Instructions For Probability Judgment Task

### Instructions

Thank you for joining our study! Please read the following instructions carefully. You will then complete a short comprehension quiz at the end before you can proceed further in the study.

##### Your task

You will evaluate stimuli representing different probabilities (p) that are presented on the screen. Your task is to determine these probabilities as accurately as possible in percentage values and type them in. For example – the probability in Figure 1 represents a probability of 0.87. Your task is to report its percentage value of 87%. The appropriate response to this would be 87. The accuracy of your typed percentages will determine your reward at the end of the experiment.

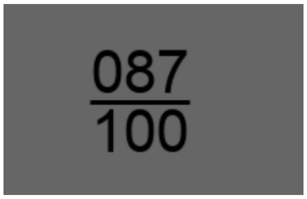


**Figure 1** Throughout the experiment, each round will present fractions representing probabilities that you have to approximate as percentages and report as accurately as possible.

##### Important details of the experiment:

#### Different probability ranges

The experiment has **two parts**, with different **ranges of probabilities**, as explained below. The order in which you do these two parts will be randomly selected. Before each part, you'll be informed of the specific range of probabilities that will be presented.

1) **Full range part**: You will be shown probabilities ranging from 0.01 to 0.99 (the full range) uniformly. The least percentage you will see in this block will be 1% while the highest will be 99%.

**Wide range of percentages between 1% and 99%**

2) **Restricted range part**: The restricted range part of the experiment itself has three sub-sections that correspond to three different restricted ranges.
   a. In the **lowest-restricted-range** sub-section, you will only be presented probabilities in the narrow range between 0.01-0.33. In these blocks, you will never see a percentage higher than 33%.

   **Percentages always between 1% and 33%** | Example probability → $\frac{17}{100}$

   b. In the **mid-restricted-range** part, you will only be presented probabilities in the narrow range between 0.35-0.65. In these blocks, you will never see a percentage lower than 35% and higher than 65%.

   **Percentages always between 35% and 65%** | Example probability → $\frac{41}{100}$

   c. In the **highest-restricted-range** part, you will only be presented probabilities in the narrow range between 0.66-0.99. In these blocks, you will never see a percentage lower than 66%.

   **Percentages always between 66% and 99%** | Example probability → $\frac{79}{100}$

#### Different complexity of probability presentation

Probabilities will be **presented with two complexity levels**:

1) In **easy blocks,** probabilities are presented as easy fractions (x/y) with a fixed y of 100 – only the x changes (see Figure 2A). The x therefore represents the probability in a manner that is easy to convert to a percentage.

2) In **hard blocks**, both the x and y in the fraction (x/y) are randomly selected 4-digit numbers that change in each round (see Figure 2B). You will thus have to estimate the probability, by approximate division of x and y. This x/y therefore represents the probability in a manner that may be harder to convert to a percentage.

a) 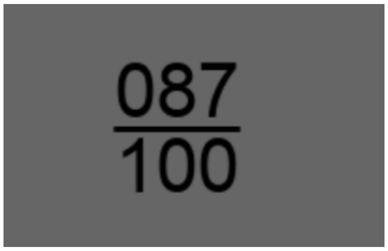

b) 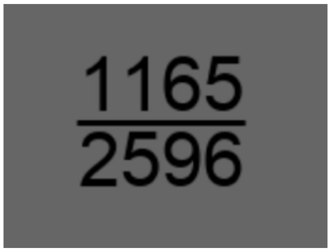


**Figure 2** a) Easy Blocks: Some of today's blocks present probabilities as fractions with a fixed denominator of 100. Only the numerator changes between rounds. b) Hard Blocks: The remaining blocks will present the probabilities as complex 4-digit fractions, with both the numerator and the denominator changing between rounds.

It is more difficult to approximate the probability/percentage in the hard blocks, but this is part of the experiment. Please try to report percentages as well as you can in both the easy and hard conditions. **You are not permitted to use any type of calculator - if you are seen using a calculator, we will have to disqualify you from the experiment without any payment.**

**Please note** -

- The order in which you will see the full and the restricted range part of the experiment is randomly determined. Within the restricted range part, whether you see the highest, middle or lowest narrow range is also determined randomly. So always pay attention at the start of a block to see which condition you are in!

- The order of “easy” and “hard” blocks within each range part is also determined randomly.

- Please pay close attention to the information provided before each block, which specifies the condition you'll encounter.

**Details on your response**

For every presented probability fraction, please input the corresponding percentage value by typing numbers at the top of the keyboard, then **press <enter> to finalize your choice.** Note that this is **necessary to record your response.** As you type, the numbers will be displayed in black; pressing <enter> will turn them green and indicate your final decision.

After 0.5 seconds of your response, the next round will start, and you will be presented with the next fraction. Once you have submitted your response, you cannot change your input anymore, so try to ensure that the percentage you entered is correct.

You have at most 5.5 seconds to indicate the percentage, ranging from 1 to 99. If you submit a number outside this range, then the answer won't be accepted and the input turns red. You can still modify your input if this happens (if your maximum time of 5.5 seconds is not over), by pressing <backspace>. You will be paid based on the accuracy over all responses you make today. The payment mechanism is designed in a way such that it will be optimal for you to be as accurate as possible within the given time in each round.

**Payment Details**

At the start of the experiment, you receive 10 CHF as a participation fee and an additional 40 CHF allocated for the experiment.

**How Penalties Work:**

- Each round, you'll see a probability as a fraction and report your estimated percentage. If your estimate is different from the actual percentage, this is a deviation from the true value.
    - If the actual probability is 0.16, this means that the actual percentage is 16% and suppose you report 20. In this scenario, your deviation is 4.
- You will be penalized based on your average percentage deviation across all rounds.
- The penalty is calculated as the average percentage deviation multiplied by 2.5. For example:
    - If your average deviation is 4 across all rounds, you will face a total penalty of 4*2.5 = 10 CHF and you will walk home with 40 CHF.

**Important:**

- Suppose you report 50% as the percentage for all rounds without considering the actual probabilities, your average deviation will be 25 per round.
- This leads to a penalty of 25*2.5 = 62.5 CHF penalty. Since you are only allocated with 40 CHF however, you'll lose the entire 40 CHF, and only take home the 10 CHF participation fee.
- If you miss a round, a deviation of 25 (maximum possible) will be used to calculate your penalty for that round, reducing your earnings.

**To Maximize Earnings:**

- Be as accurate as possible with your estimates.
- Respond promptly in every round. If you miss a round, it will reduce your earnings.

Your final payout, including the participation fee, will be displayed at the end of the experiment.

**Summary**

1. **Estimation task:** In each round, specify the percentage that you see on the screen. You have a **Time limit** of 5.5 seconds to type your answer and **submit it by pressing <enter>. You do not have to wait 5.5 seconds for the next probability if you respond earlier.**

2. This experiment has **two parts**: You are randomly assigned to complete one of them first and the other next.
    a. In one part of the experiment, you will complete **6 blocks (3 easy and 3 hard)** with probabilities covering the **full range from 1-99%.**
    b. In another part of the experiment, you will complete **6 blocks (3 easy and 3 hard)** where probabilities will be in a **restricted range**. You will randomly be assigned to complete either:
        i. The 1-33% (lowest restricted range).
        ii. Or the 35-65% (middle restricted range).
        iii. Or the 66-99% (highest restricted range).

3. We implemented a payment mechanism that makes it optimal for you to indicate the percentages in each round as accurately as possible.

4. **Important:** Always try to indicate the percentage you see in each round. If you fail to do so, or if you keep generating invalid responses (below 1 or above 99) and that round is selected, you will be penalized with a deviation of 25 bringing down your payment.

5. **Potential earnings:** These can range between 10 and 50 CHF, based on your performance.

6. **Warning:** Don't use a calculator or you will be disqualified from experiment without pay.

If you have any further questions, you can call the experimenter and clarify these. If you do not have any further questions, you can begin answering the comprehension questions on the screen. For the multiple-choice comprehension questions that follow, simply click the number 1 at the top of your keyboard if you think the right answer is option 1 and 2 at the top of your keyboard if you think the right answer is option 2.

From now on, you will only need the keyboard to complete the study. You will not need the mouse at all. Use the **number keys on the top of the keyboard**.
